# Medial Frontal Theta Reduction Impairs Rule Switching via Prediction Error

**DOI:** 10.1101/2025.06.24.661140

**Authors:** Yifei Zhang, Feng Deng, Jiajun Liao, Tom Verguts, Qi Chen

## Abstract

Cognitive flexibility, the ability to switch behavior in response to changing rules in an uncertain environment, is crucial for adaptive decision making. Prior research has hypothesized a key role of prediction error and theta oscillations in medial frontal cortex in this process. However, the causal link between such processes remains to be established. To address this, we combined neural stimulation, EEG, behavioral measurement, and computational modelling. Specifically, we applied high-definition transcranial direct current stimulation (HD-tDCS) to modulate theta oscillations as measured via EEG followed by a probabilistic reversal learning task. We find that anodal stimulation reduces theta power and rule prediction error, and it increases the number of trials needed to reliably switch between rules. These findings support the role of rule prediction error signaling as a key mechanism linking neural oscillations to behavioral adaptation and highlight the importance of theta power and rule prediction error for cognitive flexibility.

**Significance statement:** Cognitive flexibility—the ability to adjust behavior when rules change—is critical for adaptive behavior in uncertain environments. Although prediction error signaling and theta oscillations in medial frontal cortex have been proposed as key mechanisms, their causal relationship remains unclear. Here, we combine high-definition transcranial direct current stimulation (HD-tDCS), EEG, behavioral assessment, and computational modeling to establish a mechanistic link. We show that anodal stimulation reduces frontal theta power and rule-level prediction errors, leading to less efficient rule switching. These findings provide causal evidence that supports behavioral flexibility, advancing our understanding of the neural computations underlying adaptive decision making.

## Introduction

Humans flexibly shift between task rules across diverse contexts, a capacity known as cognitive flexibility (Dove et al., 2000; Cools et al., 2002; Izquierdo et al., 2017). This ability is impaired in various psychiatric disorders, such as autism, leading to persistent errors and delayed switching (Geurts et al., 2009; Peterson et al., 2009; D’Cruz et al., 2013; Crawley et al., 2020). However, the neural substrates and computational mechanisms underpinning adaptive rule switching remain unknown. Here, we combined behavioral modeling with EEG measurement and electrical stimulation to address this issue.

Prediction error—the discrepancy between expected and actual outcomes—is crucial for updating behavior when rules change (O’reilly, 2013; Schultz et al., 1997). In probabilistic reversal learning tasks, participants must adapt to dynamically changing rules by integrating feedback. In this context, rule prediction error refers to the discrepancy between current belief about the task rules and feedback that signals whether that belief is valid. Unlike stimulus - action prediction error, rule prediction error represents higher-order violations of expectations about task rules and are critical for deciding when to abandon an old rule in favor of a new one (Collins and Frank, 2013; Wilson et al., 2014). Lesion studies showed that selective lesions in the medial frontal cortex (MFC) disrupt behavioral flexibility and reduce sensitivity to negative feedback, indicating MFC’s essential role in prediction errors and adaptive behavior in non-human primates (Kennerley et al., 2006; Rudebeck et al., 2008). Recent animal study has provided causal evidence for the neural substrates underlying this process. Cole et al(2024) demonstrated in rodents that the MFC is necessary for computing rule prediction error signals: MFC inactivation directly impaired behavioral adjustment following rule switches. However, in humans, the precise relationship between rule prediction error and behavioral adjustment, along with its underlying neural mechanisms, remains to be fully elucidated.

MFC, particularly theta oscillations in this brain area, are central to processing prediction errors and guiding adaptive behavior (Botvinick et al., 2004; Ridderinkhof et al., 2004; Mansouri et al., 2006; Badre et al., 2010; Kolling et al., 2012; O’reilly, 2013; Cavanagh and Frank, 2014; Donoso et al., 2014). Theta power increases following negative feedback, presumably supporting rule updating. This in turn informs decision-making and behavior adaptation, thereby supporting flexible rule-switching. Here, the Sync model proposed by Verbeke and Verguts (2021) provides a framework for understanding how MFC theta oscillations interact with rule prediction error signals. The Sync model has a hierarchical structure, using rule prediction errors to monitor environmental shifts in which task is currently relevant, and stimulus - action prediction errors to update stimulus processing in task-specific modules (Collins and Frank, 2013; Badre and Nee, 2018). When the accumulated rule prediction errors reach a certain threshold, theta amplitude increases, which facilitates the switch to a new task rule module. This mechanism has been validated computationally and is supported by neurophysiological (EEG) evidence. However, a critical gap remains in understanding the causal role of medial frontal theta oscillations in modulating (rule) prediction error computations that underlie such processes.

To test causality, here we applied high-definition transcranial direct current stimulation (HD-tDCS) to MFC. HD-tDCS allows for non-invasive modulation of cortical excitability (Nitsche et al., 2008; Dennison et al., 2019) and has been used to alter cognitive control and social learning (Reinhart and Woodman, 2014; Liu et al., 2023; Qu et al., 2024).

Building on prior research linking midfrontal theta activity to prediction error processing, we hypothesized a causal pathway in which theta oscillations influence behavioral adaptation via rule prediction error. We further predict that active stimulation of the MFC will causally modulate behavioral adjustment by altering theta-related prediction error signaling. Our findings provide insights into the neural mechanisms of cognitive flexibility and may inform interventions for disorders involving impaired adaptive behavior.

## Materials and Methods

### Participants

The study recruited 50 right-handed individuals with either normal or corrected-to-normal vision. Before participating, individuals signed informed consent. The experimental procedures received approval from the Ethics Committee of the School of Psychology at South China Normal University (Approval No. SCNU-PSY-2021-042). After excluding two participants whose accuracy was more than two standard deviations below the overall mean accuracy, the final sample consisted of 48 participants (18 females; 30 males; age range: 19–28 years, *M* = 22.40, *SD* = 1.95).

### Apparatus and Materials

The experiment used PsychoPy software (Peirce et al., 2019) for implementation. Electrophysiological recordings were gathered using a Brain Products system with 64 Ag/AgCl electrodes, following the standard international 10-20 configuration, and EEG data were sampled at 1000 Hz. Initially, FCz was used as reference electrode, and these were later re-referred to the left and right mastoids during the preprocessing stage.

Behavioral data analysis was carried out using R software (R Core Team, 2017), and computational models were estimated with the differential evolution method within the SciPy package (version 1.4.1) for Python (version 3.7.6). Preprocessing of electrophysiological data was completed in MATLAB R2023b, using the EEGLAB pipeline (Delorme and Makeig, 2004).

### Code and data accessibility

All code used to produce the results reported in this paper is publicly available at https://github.com/yifei0710/tDCS-EEG. Data will be made available upon reasonable request.

### Procedure

All experiments began with screening questionnaires, then brain stimulation, and were followed by an EEG recording while subjects carried out the probabilistic reversal learning task (Figure 1A). Each subject participated in three different stimulation conditions (i.e., anode, cathode, and sham) on different days with order counterbalanced across subjects (Figure 1B). The time interval between testing days was 24 hours which avoids ordering confounds related to repeated brain stimulation exposure (Xu et al., 2023).

**Figure 1.**
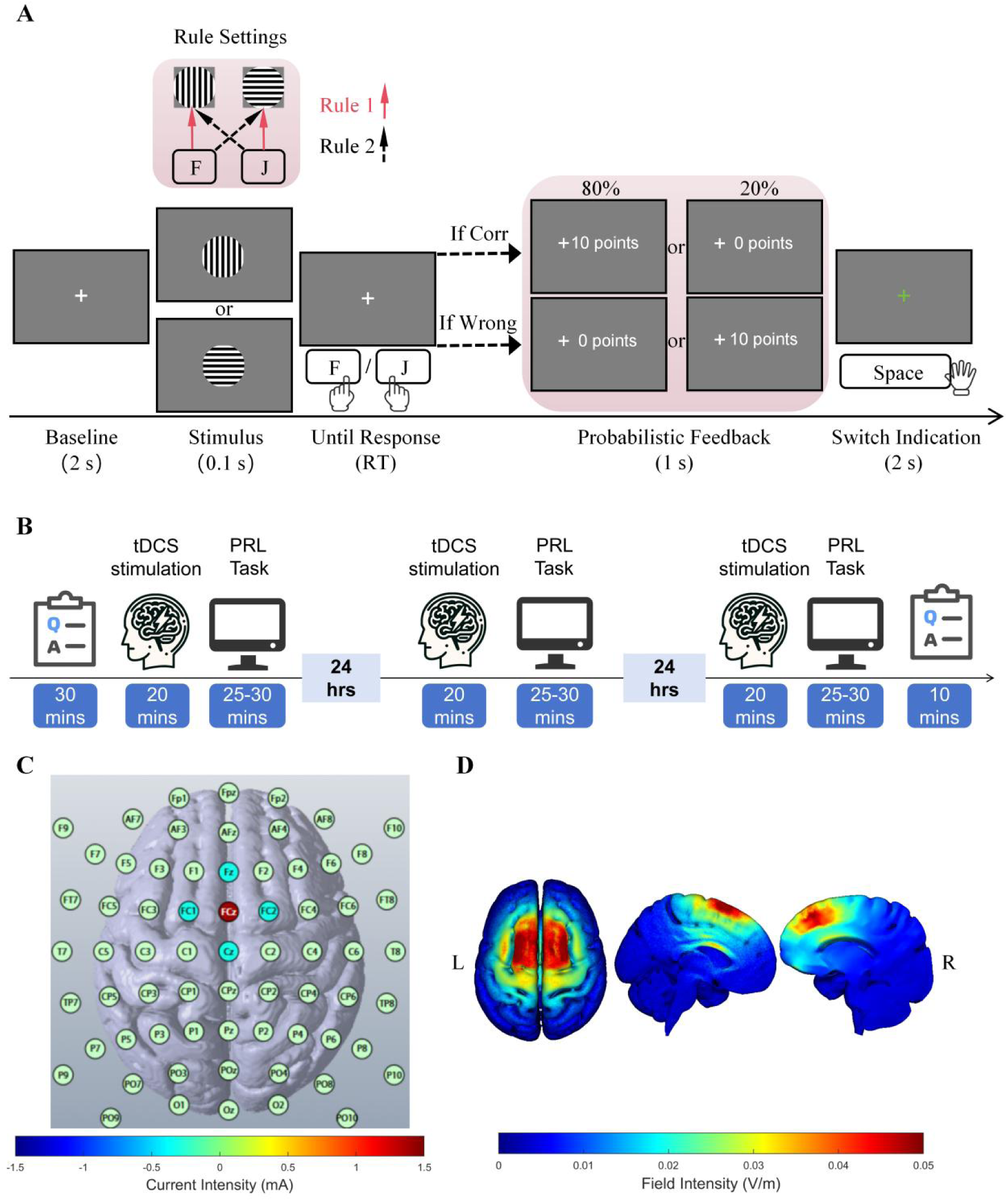
(A) Probabilistic reversal learning (PRL) task. Each trial started with a central fixation cross presented for 2 seconds, followed by the display of one of two stimuli (vertical or horizontal grating) for 100 ms. The screen remained a white fixation cross until a response was made. Probabilistic feedback was then presented at the center of the screen. Following the feedback, a green fixation cross appeared, and participants were instructed to press the space bar if they inferred a rule change. (B) The experiment was conducted across three separate days, with a minimum interval of 24 hours between sessions. Each session began with a 30-minute preparation and screening questionnaire, 20-minute HD-tDCS stimulation, followed by the task and a debriefing at the end of experiment. (C) The MFC neuromodulation protocol. HD-tDCS was administered using a 5-electrode montage targeting the medial frontal cortex (MFC). The electrode was placed at FCz, surrounded by four return electrodes at Fz, FC1, FC2, and Cz, following the international 10–20 system. (D) The current-flow models on 3D reconstructions of the cortical surface.

#### Experimental design

The experimental task paradigm was adapted based on Verbeke et. al (2021). During this task (Figure 1A), subjects had to learn two distinct stimulus - action mappings that were intermittently reversed throughout the experiment. Each trial began with a white fixation cross at the center for 2000 ms, which was followed by a stimulus display for 100 ms. The stimulus consisted of a circular grating framed by a raised-cosine mask spanning 7 visual degrees, oriented either vertically or horizontally. After seeing the stimulus, the screen turned into a white fixation cross until participants made their response by pressing the ‘F’ (left) or ‘J’ (right) key on a qwerty keyboard. In Rule 1, a vertical orientation solicited a left response, while a horizontal orientation required a right response; these configurations were reversed in Rule 2. The experiment included 240 trials with seven rule reversals occurring randomly, each drawn from a uniform distribution between 15 and 45 trials after the previous reversal. Feedback followed each response, indicating ‘+10 points’ for rewarded outcomes, ‘+0 points’ for unrewarded ones, or ‘Respond faster!’ if response times (RTs) exceeded 1000 ms. Feedback was probabilistic, with the probability of reward feedback being 80% for correct responses and 20% for incorrect responses. Following feedback, a green fixation cross reappeared for another 2000 ms, instructing participants to press the space bar upon detecting a rule switch. This setup aimed to determine each participant’s personal ‘switch threshold,’ based on the Sync Model. However, as these self-reports were not the focus of the current, they were not further analyzed in this study. Participants were allowed short breaks around every 60 trials; no rule switch occurred within the 10 trials leading up to the break.

#### HD-tDCS

The high-definition transcranial direct current stimulation (HD-tDCS) was administered using a battery-powered, constant current device together with a 5-channel high-definition transcranial electrical-current stimulator (Soterix Medical). HD-tDCS targeted the MFC to influence neural activity (Figure 1C). The stimulation lasted for 20 minutes at 1.5 mA and was centered on the FCz electrode as per the International 10–20 system. Five electrodes were placed at Fz (−0.375 mA), FC1 (−0.375 mA), FCz (1.5 mA), FC2 (−0.375 mA), and Cz (−0.375 mA), where FCz was the active electrode, and the others were return electrodes (Reinhart and Woodman, 2014). The FCz placement was determined by current flow models (Figure 1D), which suggested this location effectively targets medial–frontal structures linked to producing medial–frontal negativities associated with prediction error signaling (Gehring and Willoughby, 2002; Reinhart et al., 2015). Studies on prefrontal cortex stimulation have found that the behavioral and neural effects of anodal tDCS can persist for approximately 5 hours post-stimulation (Reinhart and Woodman, 2014). This duration exceeds the length of our experimental sessions (approximately 30 minutes), ensuring that the stimulation effects were maintained throughout the task. A sham condition was also utilized, employing the same electrode setup and procedures. In the sham condition, current gradually increasing and decreasing at the start, and the end of the 20-minute duration to simulate the sensation of receiving active stimulation. Participants were kept unaware of whether they received active or sham stimulation, and after debriefing, it was confirmed that none were aware of the stimulation polarity. Furthermore, aside from mild tingling under the electrodes, no adverse effects were reported.

### Statistical Analysis

#### Behavioral data analysis

We define *perseverative errors* as trials in which participants persist with the previously valid rule and fail to switch to the new correct rule after a reversal (Weiss et al., 2021). *Regressive errors* are defined as instances where a participant, after initially switching to the correct rule, subsequently reverts to the previously incorrect one. Additionally, we introduce the measure *trials to stable rule switch*: if a participant switches to the correct rule and maintains correct responses for the following two trials, we consider a stable switch to be completed. We then calculate the number of trials from rule reversal to stable switch. This measure includes the stable switch trial itself. Trials to stable rule switch offers a direct and integrative measure of learning stability, and it will serve as a key measure in subsequent modeling and EEG analyses.

#### Computational Modeling

##### Sync Model

The Sync Model (Figure 2) provides a mechanistic account of how MFC theta oscillations contribute to cognitive flexibility during probabilistic reversal learning. It is fully reported in Verbeke et al (2021) but we repeat it here for completeness. The model consists of two interconnected and hierarchically related modules: the (lower level) Mapping Unit and the (higher level) Rule Unit. The Mapping unit represents stimulus-action associations. Specific stimulus features (e.g., striped patterns) are mapped onto corresponding motor responses (e.g., pressing “F” or “J”). These mappings are updated based on feedback, which reflects whether the chosen response was correct or incorrect. The Rule unit monitors feedback to regulate the Mapping unit. It comprises lateral frontal cortex (LFC) and medial frontal cortex (MFC) components. The LFC encodes different task rules (e.g., Rule 1 vs. Rule 2), while the MFC tracks rule prediction errors generated when feedback indicates that the current rule may no longer be valid. The MFC communicates with the LFC via theta oscillations. When feedback indicates that the current rule is invalid, a rule prediction error is computed. An accumulation of rule prediction errors triggers MFC theta bursts, which itself can trigger a rule update in the LFC. This mechanism allows the system to switch from one rule module (e.g., Rule 1) to another (e.g., Rule 2) adaptively. Feedback from the Mapping unit informs the Rule unit whether current stimulus-action mappings are successful. The rule prediction errror 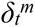 for rule *m* is defined as a discrepancy between the Reward *R_t_* and the expected value 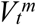 for rule *m* at trial *t* :

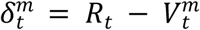

and 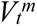 is updated under the currently active rule module as follows:

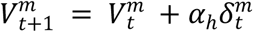

**Figure 2.**
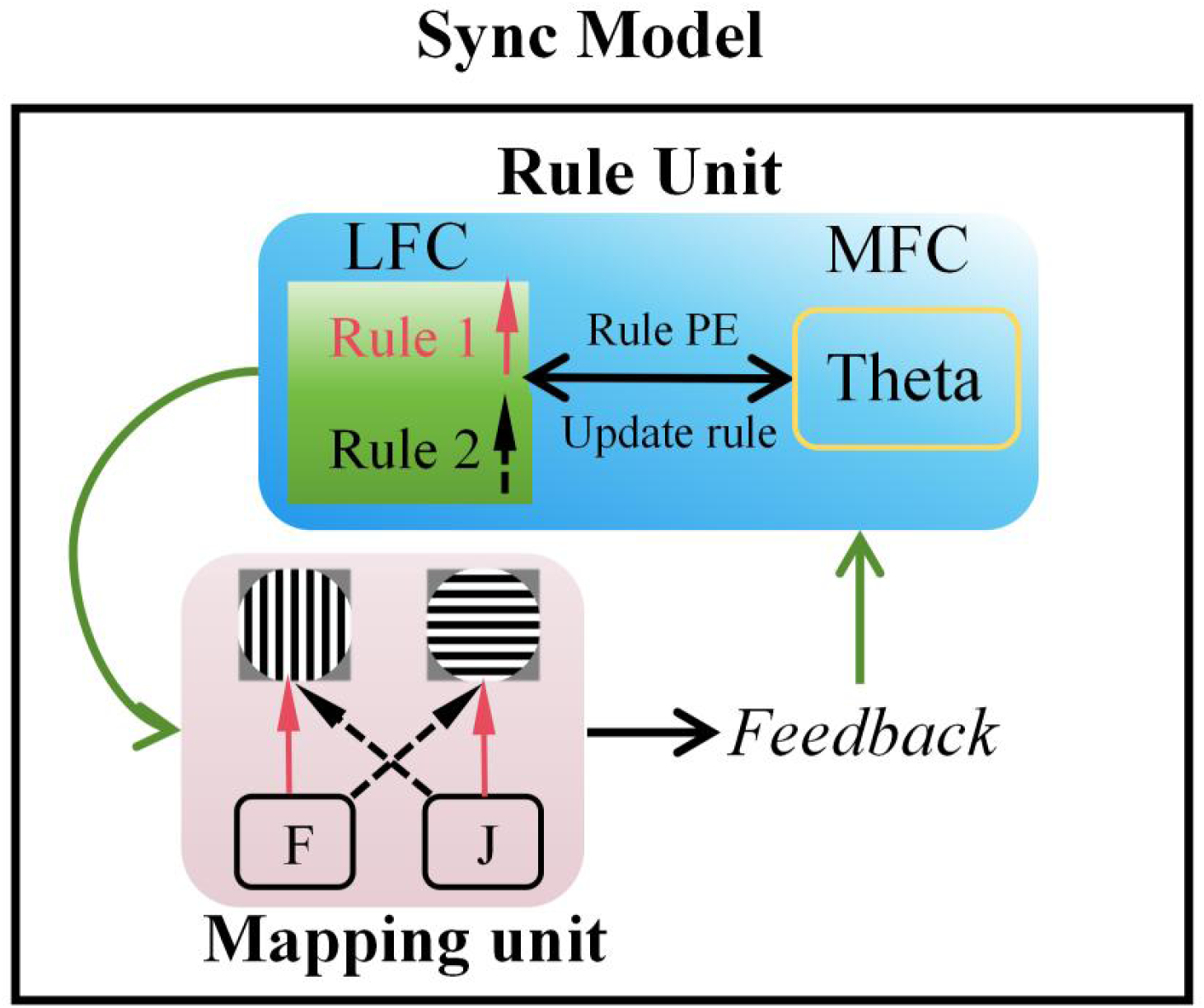
Schematic overview of the sync model. The Sync model explains flexible rules switching through the interaction between medial frontal cortex (MFC) and lateral frontal cortex (LFC). The Mapping Unit encodes stimulus-action mappings (e.g., grating orientation mapped to keys F or J), while the Rule Unit monitors rule context and controls switching. The LFC maintains the currently active rule (e.g., Rule 1 or Rule 2), and the MFC generates theta oscillations that encode rule prediction error (Rule PE) when feedback indicates a mismatch between the expected and actual outcome. Upon sufficient accumulation of Rule PE, the MFC signals the need to update the rule, shifting control from one rule module to the other. This hierarchical coordination enables dynamic adjustment of behavior based on environmental feedback.

The parameter *α_h_* refers to the hierarchically higher Rule unit learning rate. The difference between 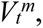 and actual observed reward *R*_t_ (i.e., the rule prediction error) is summed in Accumulator neuron (*A*) as shown below:

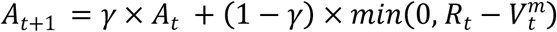

Since switches are only required in the presence of negative feedback, the Accumulator neuron is selective for negative prediction errors. That is, only negative rule prediction errors accumulate. The Cumulation parameter *γ* determines the contribution of a single prediction error on the Accumulator neuron. A low value of the Cumulation parameter leads to a stronger weight on individual rule prediction error, causing the system to frequently switch between rule modules. The model switches to a new rule module in the Mapping unit when the Accumulator neuron value exceeds a rule switch threshold of 0.5.

In addition to the Sync model, we estimated two baseline reinforcement learning models for comparison. First, the classical Rescorla-Wagner model (Wagner and Rescorla, 1972) updates the expected values of stimulus-action mappings using a fixed learning rate based on trial-wise outcome prediction errors. This model assumes a constant sensitivity to prediction errors across trials. Second, the Augmented Learning Rate model (Bai et al., 2014) builds on the Rescorla-Wagner model by introducing a variable learning rate that changes with the magnitude of prediction error. This allows the model to adjust learning rates dynamically, offering improved flexibility in non-stationary environments.

##### Model Fitting and Evaluation

We fitted all three models with behavioral data using the differential evolution optimization method available in the SciPy package for Python. For fitting behavioral data, the Sync model was simplified (Verbeke and Verguts, 2019). This simplified version of the model introduces a hard gating process between rule modules and a softmax response selection mechanism, providing an effective means of modeling behavioral data. Within a rule module, an action is chosen as follows:

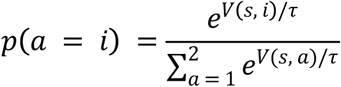

where *p* (*a = i*) represents the probability of selecting action *a = i* in a given stimulus *s*. *V* (*s, i*) is the value associated with a specific stimulus - action pair (*s, i*); it reflects the expected reward or utility of performing action *i* to stimulus *s*. The temperature parameter *τ* controls the degree of exploration versus exploitation in the decision-making process. The denominator involves a summation over all possible (in this case, 2) actions to normalize the probability distribution.

#### Electrophysiological data analysis

##### Preprocessing

Prepossessing included filtering, interpolation, re-referencing, artifact rejection, independent component analysis (ICA), epoching, removing bad epochs, and downsampling. This was performed using EEGLAB in MATLAB. EEG data were offline re-referenced to the average of the mastoid electrodes. Segments including breaks and other offline intervals were manually removed from the analysis. Noisy electrodes were identified; data from surrounding electrodes were used for interpolation at all time points. Specifically, interpolation was performed on one electrode for one participant and on two electrodes for two other participants. Data was bandpass filtered between 1 and 48 Hz to remove slow drifts and at 50 Hz line noise. Eye blinks and motor-related artifacts were eliminated through ICA in EEGLAB. After ICA, the data were divided into epochs time-locked to feedback and stimulus onset. Baseline activity, extracted from epochs locked to stimulus onset, covered −1500 ms to −500 ms relative to stimulus onset and was subtracted from all epochs. Removing bad epochs, an average of 7.5% of epochs was excluded based on an amplitude threshold of ±100 µV and an improbability test, defined as 6 standard deviations for individual electrodes and 2 for all electrodes.

##### Time-frequency analysis

Time-frequency analyses explored theta band (4–8 Hz) power dynamics. Before conducting time-frequency analyses, the data were down sampled to 500 Hz. Utilizing code adapted from Cohen (2014), time-frequency decomposition was executed. The procedure employed complex Morlet wavelets to explore frequencies from 2 to 48 Hz, divided into 25 steps spaced logarithmically. The number of cycles for each frequency ranged from 3 to 8, also segmented into 25 logarithmic steps.

##### Power Computation

Baseline correction involved normalizing the power estimates for each subject, electrode, and frequency against the average baseline activity measured from -1500 ms to -500 ms before the stimulus onset during the baseline period across all 240 trials. Following baseline correction, power values were transformed into decibels (dB) to enable condition comparisons. Trials with delayed responses were excluded before the final analyses to maintain data quality and reliability.

We calculated a contrast between z-scored power for trials yielding negative feedback and those with positive feedback (Z_Neg-Pos_). A nonparametric clustering method (Maris and Oostenveld, 2007), was then applied to these contrast values. The statistical value distribution was generated, and the most extreme 1% on both sides (for a two-sided test) was selected for clustering. Clusters were formed by grouping adjacent time points, frequencies, and channels that surpassed the threshold. The cluster-level statistic was determined by multiplying the number of elements in each cluster by the highest statistical value in that cluster. A 5% significance threshold was set using a nonparametric permutation test with 1000 iterations. Clusters passing this test were used in further analyses, ensuring reliable identification of significant power differences associated with feedback processing.

##### Time-Frequency Comparison Across TDCS Stimulation Conditions

We aim to contrast the theta cluster activity differences between the sham and active conditions. Based on the variable Z_Neg-Pos_ from the time-frequency analysis, we subtracted the anodal condition from the sham condition to obtain the contrast values. The same nonparametric statistical method as previously described was applied to correct for multiple comparisons.

##### Pre-Stimulus Baseline Comparison Across TDCS Stimulation Conditions

To assess whether tDCS modulated ongoing neural activity independent of feedback-related responses, we examined pre-stimulus theta power. Baseline theta power was defined as the mean power within the theta band (4–8 Hz) from −500 to 0 ms before stimulus onset during the baseline period. First, we computed the average theta power for all trials and frequency bands for each electrode, thereby creating topographical maps of baseline activity for all stimulation conditions. Then, we subtracted the baseline power of the sham condition from the baseline power of the active condition to reveal spatial patterns of differential activity. By focusing on prefrontal regions, we selected electrodes corresponding to medial–frontal structures (i.e., FCz, FC1, FC2, Fz, F1, F2, AFz, AF3, AF4) and computed the average baseline theta power across these sites.

#### Mediation Analysis

To assess the mediation effect of rule prediction error between theta and trials to stable rule switch, we conducted a mediation analysis under both the sham and active conditions. The (trial by trial) mediation analysis used only the rule prediction error and theta power measured on the trials in which the rule reversal took place.

In this mediation analysis, theta power served as the independent variable, rule prediction error as the mediator, and the number of trials required to achieve a stable rule switch as the dependent variable. The indirect effect was assessed through the product of the coefficients for the paths from theta to rule prediction error (*a* path) and from rule prediction error to trials to stable rule switch (*b* path). The direct effect of rule switch theta on trials to stable rule switch (*c*’ path) was also computed. The significance of the indirect effect was tested using bootstrapping methods with 5,000 resamples, providing confidence intervals for the mediation paths. The results were evaluated separately for each stimulation condition to determine whether the mediation effect was condition-dependent.

#### Correlation Analysis on EEG data in Stable Rule Switch Contrast Across TDCS Stimulation Conditions

To investigate the relationship between EEG-based constraints in feedback processing and observed behavioral deficits, we conducted Pearson correlation analyses. Specifically, we assessed the correlation between the differences in pre-stimulus baseline theta power (sham vs. active) and the differences in theta power during feedback processing with the differences in trials to stable rule switch. This analysis aimed to determine whether the baseline theta activity and the feedback theta activity contribute to cognitive flexibility. Correlation coefficients were calculated using R software (R Core Team, 2017), and significance was determined using two-tailed tests with an alpha level of 0.05.

## Results

### Behavioral data: Accuracy and Reaction Time

Accuracy and reaction time on the probabilistic reversal learning task were examined using repeated-measures ANOVA across the sham, anodal, and cathodal conditions. The analysis revealed no significant effect of stimulation on overall accuracy (*F* _(2, 93.11)_ = 0.84, *p* = .44), nor on reaction time (RT) (*F* _(2, 93.64)_ = 0.10, *p* = .91). Specifically, mean accuracy in anodal (*M* = 79.73, *SD* = 4.31) or cathodal (*M* = 79.66, *SD* = 5.51) conditions did not significantly differ from that in sham condition (*M* = 80.80, *SD* = 4.76), as shown in Figure 3A. Likewise, mean RT in the sham condition (*M* = 542.51 ms, *SD* = 62.20) was comparable to that in the anodal (*M* = 545.82 ms, *SD* = 61.10) and cathodal (*M* = 540.63 ms, *SD* = 54.30) conditions (Figure 3B). These findings suggest that tDCS stimulation did not produce any detectable changes in accuracy and RT.

**Figure 3.**
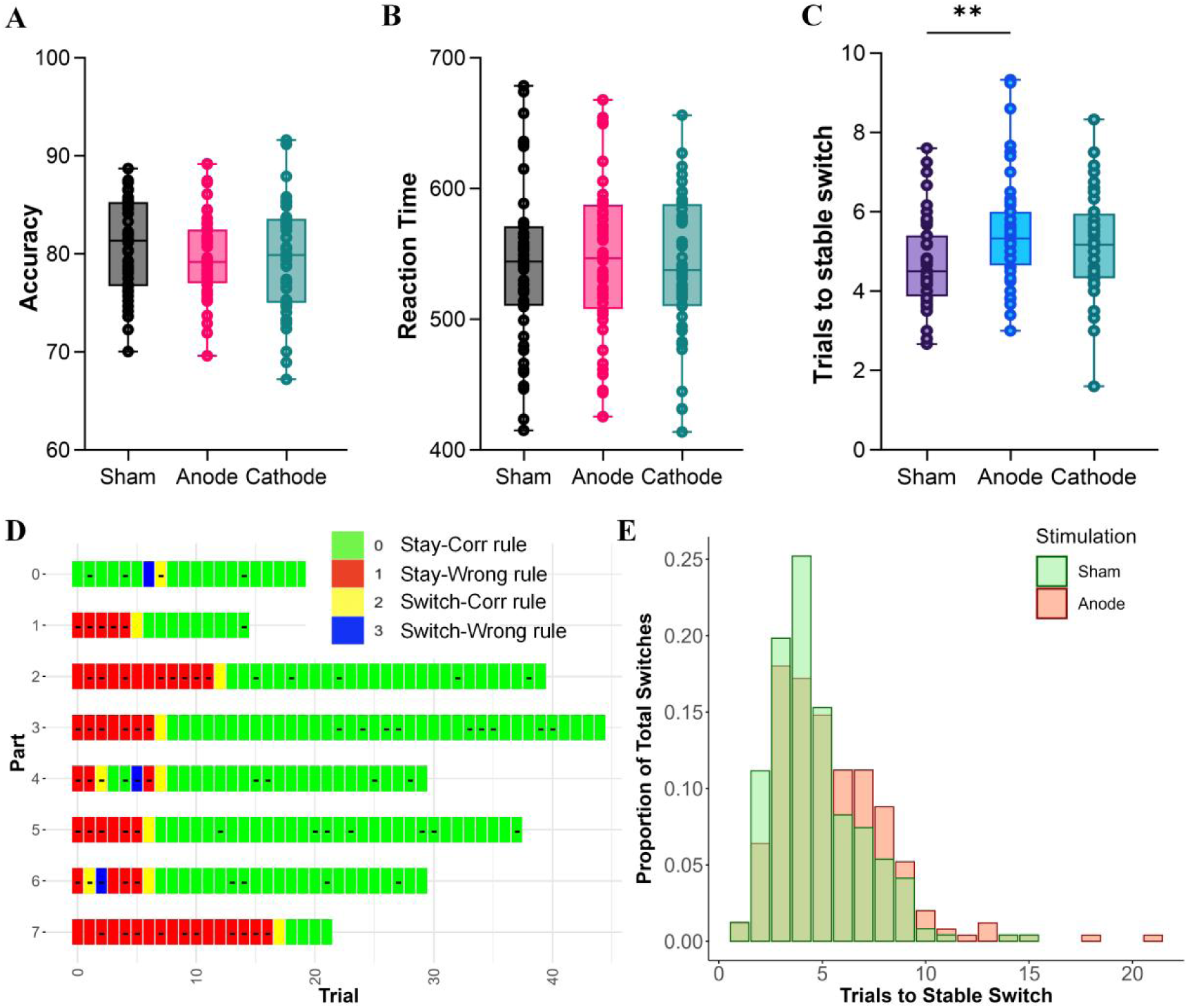
Behavioral performance across stimulation conditions. (A) Accuracy under sham, anodal, and cathodal stimulation conditions. (B) Reaction time under the same stimulation conditions. No significant differences were observed across conditions in either accuracy or reaction time. (C) Number of trials required to achieve stable rule switch, with a significant increase in the anodal condition compared to sham. Box plots represent the distribution of individual data points, with the central line indicating the median and whiskers showing the data range. *p < 0.05; **p < 0.01; ***p < 0.001. (D) Example of one participant’s color-coded behavior for all trials. (E) Histogram of number of trials to stable rule switch for sham and anodal conditions.

### Anodal Stimulation Delays Stable Rule-Switching

Our objective was to examine how individuals adjust their behavior in response to probabilistic feedback. As shown in one illustrated example from one participant (color-coded in Figure 3D), each rule reversal was aligned to the zero point on the horizontal axis, and the vertical axis (0–7) denotes consecutive segments, with Part 0 serving as the initial rule-learning phase and Parts 1 to 7 referring to the rule reversal learning phase. This color-coded representation was designed to visually illustrate how participants responded to negative feedback and when they chose to alter their rules based on their actual behavior adjustment. We used a four-color scheme (green for persisting with the correct rule, red for persisting with an incorrect rule, yellow for switching to the correct rule, and blue for switching to an incorrect rule) to identify three indices of rule switching responses. A “-” symbol indicates that the participant received negative feedback on that trial. By aligning reversals and mapping choice behavior trial by trial, this visualization offers an intuitive way to capture the temporal dynamics of rule adaptation and supports the validation of the three behavioral metrics.

We can thus illustrate the three fine-grained metrics defined in Methods. First, *perseverative errors* were quantified as the number of errors (trials) after each rule reversal until the participant’s first successful switch to the correct rule, including the successful switch itself. In Figure 3D, Part 6 yielded perseverative errors of 1 because one error were made before switching to the correct rule. The corresponding value for this part is coded as 2, including the successful switch trial. A repeated measures ANOVA confirmed that there were no significant differences of perseverative errors across three stimulation conditions (*F*_(2, 93.98)_ = 0.51, *p* = 0.60; sham: *M* =4.13, *SD* = 1.07; anode: *M* = 4.34, *SD* = 1.06; cathode: *M* = 4.23, *SD* = 1.19) (Supplementary Figure 2A). We also examined *regression errors*, which refer to instances where participants initially switched to the correct rule but subsequently reverted to the previous one (Supplementary Figure 2B).

Second, we examined regression errors (Supplementary Figure 2B), which refer to instances where participants initially switched to the correct rule but subsequently reverted to the previous one. Continuing with the same the example, part 6 showed 4 regression errors. Compared to the sham condition, participants in the anodal stimulation group exhibited a significantly higher number of regression errors, suggesting a greater tendency to persist with the previous rule despite changes in task contingencies (*F*_(1.88, 88.45)_ = 7.23, *p* = 0.0082; sham: *M* = 0.57, *SD* =0.78; anode: *M* = 1.17, *SD* = 1.21; cathode: *M* = 0.94, *SD* = 1.02). Post hoc analyses revealed a significant difference in regression errors between the sham and anodal stimulation conditions (*t*_(47)_ = –3.29, *p* = 0.005), with participants exhibiting larger errors under the anodal condition compared to the sham condition (mean difference = –0.600, *SE* = 0.182). The comparison between sham and cathodal stimulation was not significant (*t*_(47)_ = –2.22, *p* = 0.078), while no significant difference was found between anodal and cathodal conditions (*t*_(47)_ = 1.08, *p* = 0.533). This suggests that participants under anodal stimulation experienced more difficulty in consistently maintaining the correct rule following each reversal.

Third, a stable rule switch response requires the participant to maintain two additional correct responses following the correct switch, resulting in a value of 7 in the same example (Figure 3D). Compared to sham, the anodal condition produced significantly more trials until stable rule switches (*F*_(1.880, 88.35)_ = 5.233, *p* = 0.001; sham: *M* = 4.70, *SD* = 1.13; anode: *M* = 5.50, *SD* = 1.38; cathode: *M* = 5.17, *SD* = 1.27) (Figure 3C). Post-hoc pairwise Tukey tests revealed that participants under anodal stimulation required significantly more trials to reach stable rule switching compared to sham (*t*_(47)_ = -3.52, *p* = 0.003, Cohen’s d = -0.57). The difference between anodal and cathodal stimulation was not significant (*t*_(47)_ = 1.81, *p* = 0.178), nor was the difference between sham and cathode (*t*_(47)_ = -2.12, *p* = 0.096). An increase in trials to stable switch after anodal stimulation with a similar number of perseverative errors indicates that participants continue to make errors even after their initial switch to the correct rule, suggesting that they fail to fully utilize feedback to stabilize learning. Since significance was only found between sham and anode, Figure 3E shows the distribution of trials to stable rule switch under sham and anodal conditions. The tendency to persist with the old rule and delay the adoption of the new rule highlights a notable behavioral effect of anodal stimulation.

### Anodal Stimulation Selectively Blunts Rule Prediction Error

The Sync Model consistently outperformed both the Rescorla-Wagner and adaptive learning rate models in all model fit evaluation metrics (see Supplementary Table 1), showing that it can capture rule-switching behavior in the probabilistic reversal learning task. Hence, all subsequent analysis will be based on the Sync model.

No significant differences were observed across stimulation groups in the fitted mapping learning rate, switch learning rate, temperature, or cumulation parameters (Supplementary Figure 3A–D).

To generate simulated behavioral data (in particular, rule prediction errors), the Sync model was run trial-by-trial using the actual sequence of stimuli and feedback and using each participant’s best-fitting parameters. Two linear mixed-effects models were fitted to examine the effect of stimulation on negative and positive rule prediction error respectively. The model included stimulation as a fixed effect and a random intercept for subject to account for repeated measures. The effect of stimulation was significant for both negative and positive rule prediction error (negative: *F*_(2, 6807)_ = 13.5, *p* < .001; positive: *F*_(2, 13597)_ = 38.9, *p* < .001). For negative rule prediction error, post-hoc comparisons showed that the anodal condition significantly decreases the absolute value of rule prediction error, compared to sham (estimate = -0.021, *SE* = 0.004, *t*_(6809.4)_ = -4.84, *p* < .001, Figure 4A). The cathodal condition did not significantly differ from sham (*t*_(6804)_ = 0.872, *p* = 0.383). For positive rule prediction error, post-hoc comparisons showed that the anodal condition significantly increases rule prediction error compared to sham (estimate = 0.025, *SE* = 0.003, *t*_(13497.3)_ = 8.049, *p* < .001, Figure 4B, whereas the cathodal condition did not significantly differ from sham (*t*_(13597.2)_ = 0.889, *p* = 0.374).

**Figure 4.**
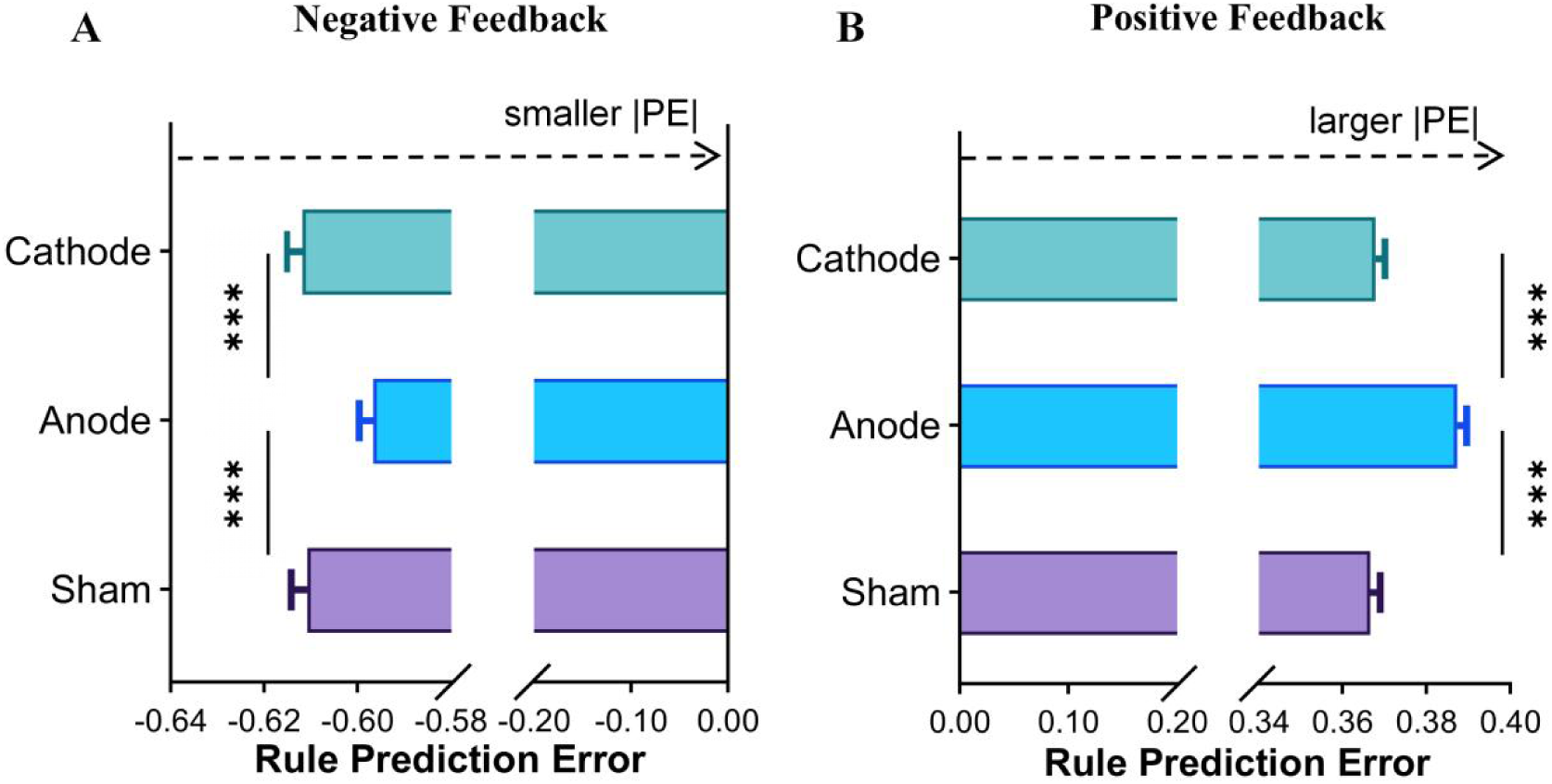
Rule prediction error comparison among tDCS conditions. (A) Negative feedback showed significant blunted rule prediction error under anodal condtion compared to the sham and cathodal conditions. (B) Positive feedback showed anodal condition significantly increased the rule prediction error compared to the sham and cathodal conditions. Error bars represent standard errors of the mean (SEM). Statistical significance is indicated as \**p* < 0.05; \*\**p* < 0.01; \*\*\**p* < 0.001.

Recall that negative feedback is critical in this experimental design to switch adaptively. To further dissociate hierarchical levels of prediction error processing, we conducted separate control analysis focusing on negative low-level prediction errors (stimulus - action prediction errors). A linear mixed-effect model was fitted for negative stimulus - action prediction errors with stimulation as a fixed effect and subject as a random intercept. The results showed no significant effect of stimulation (Supplementary Figure 4, *F*_(1,6834)_ = 0.476, *p* = 0.621). These findings suggest that anodal stimulation selectively blunted (negative) rule prediction error, not stimulus-action prediction errors.

### Anodal Stimulation Reduces Midfrontal Theta Power After Feedback

Since the behavioral and modeling results revealed significant differences only between the anodal and sham conditions, the following EEG analyses focus exclusively on these two conditions. For completeness, results related to the cathodal condition are provided in the Supplementary Materials (Supplementary Figure 5, 6). We next examined the EEG data to investigate neural dynamics at the time of feedback onset, aiming to understand how the brain responds to feedback and whether these neural responses differ between anodal and sham stimulation conditions. Cluster analysis of post-feedback power (0–1000 ms) identified two significant clusters associated with feedback processing under sham and anodal stimulation conditions (Figure 5A–D). These two significant clusters are theta cluster (4-8 Hz) and alpha cluster (8-20 Hz). Of particular interest in the context of rule-switching was a midfrontal theta cluster observed between 0 and 600 ms (Figure 5A, C). Notably, this theta cluster exhibited greater power in response to negative feedback compared to positive feedback. The theta clusters were consistently localized to midfrontal electrodes in both sham and anodal conditions (Figure 5B, D). Upon comparing the two stimulation conditions, we observed a significant reduction in theta power in the midfrontal cluster under anodal stimulation relative to sham. (Figure 5E and Figure 5F).

**Figure 5.**
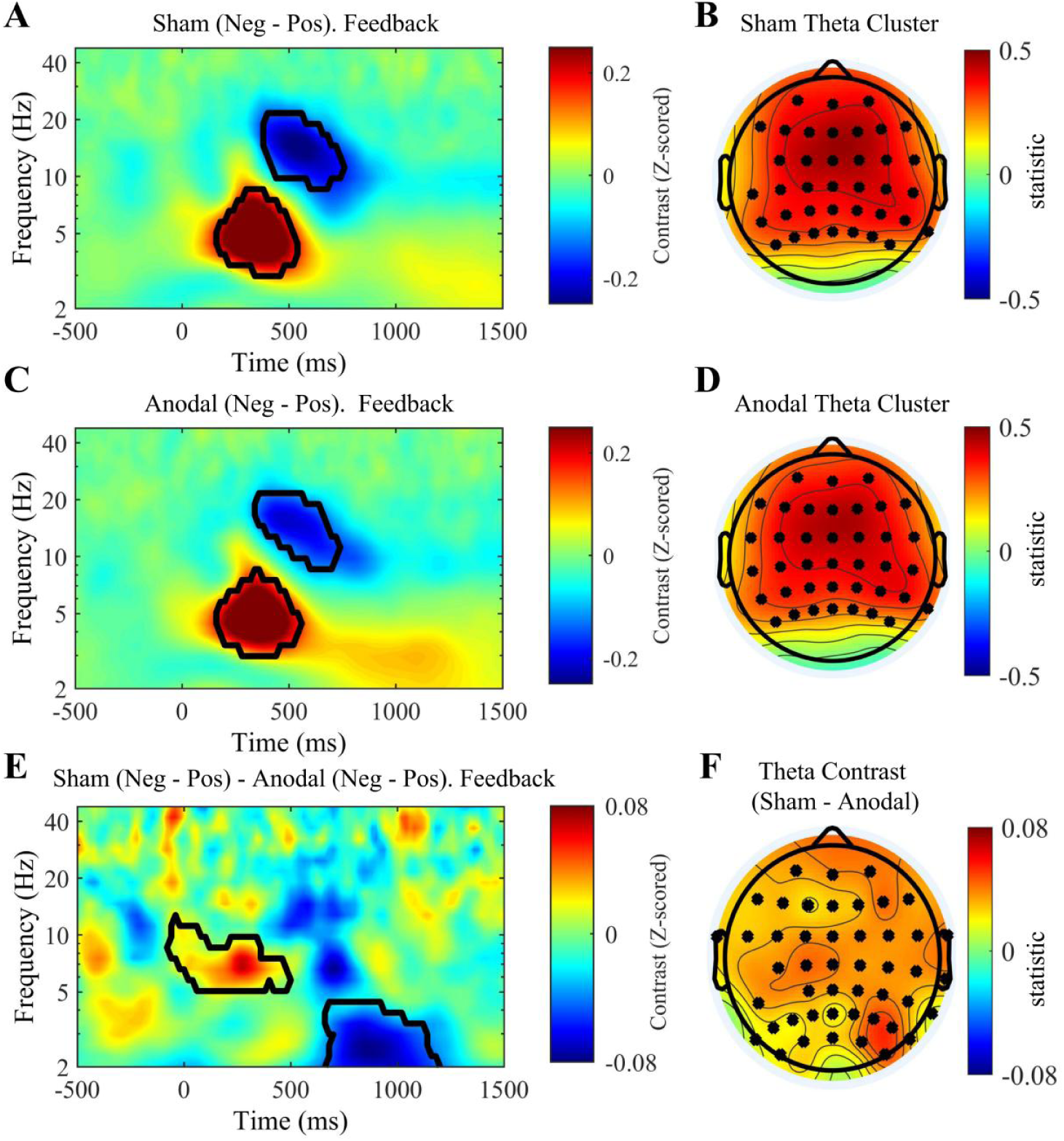
Time-frequency representations and topographic distributions of theta-band activity across stimulation conditions. (A) Time-frequency plot of the contrast between negative and positive feedback in the sham condition, with significant clusters outlined in black. (B) Topographical distribution of the theta cluster identified in the sham condition. (C) Time-frequency plot of the contrast between negative and positive feedback in the anodal condition, with significant clusters outlined. (D) Topographical distribution of the theta cluster in the anodal condition. (E) Difference in time-frequency representations between sham (negative - positive feedback) and anodal (negative - positive feedback) conditions, with significant clusters outlined. (F) Topographical representation of the Theta Neg-Pos contrast (sham - anode) from Figure E. Color bars indicate contrast values in z-scores (A, C, E) or statistical power (B, D, F).

To investigate if this reduction effect is consistent across subjects, we next extracted subject-specific Negative–Positive theta contrasts from the significant cluster identified at the group level. This allowed us to assess whether the modulatory effect of anodal stimulation varies across individuals (see Supplementary Figure 7). A one-sample t-test confirmed that the Negative–Positive theta difference between sham and anodal conditions was significantly different from zero at the participant level (*t*_(47)_ = 6.03, *p* < .001, *M* = 0.209, *SD* = 0.241), indicating a reliable reduction in theta power following anodal stimulation. Notably, 40 participants (approximately 83.3% of the sample) aligned with this pattern, suggesting that the suppressive effect of anodal stimulation on feedback-related theta activity is consistent across most individuals.

### Mid frontal theta predicts rule prediction error

We next predicted trial by trial theta power based on rule prediction errors estimated from the Rule unit (see Methods). A regression model that included the interaction between rule prediction errors and reward provided a significantly better fit than a model with rule prediction errors as the sole regressor (χ^2^(1, *N*=48) = 110.27, *R*^2^ = 0.203, p < .001). As illustrated in Figure 6A, B, the results showed a robust negative linear relationship between theta power and negative rule prediction errors. These findings suggest that theta power selectively increases in response to negative rule prediction errors, consistent with the Sync model’s proposal that frontal theta reflects the accumulation of such errors to guide rule switching. The analysis revealed a significant main effect of rule prediction error (*F*_(1,13816)_ = 56.34, *p* < .001, *β* = -0.403) and a significant interaction of rule prediction error and reward (*F*_(1,13780)_ = 62.90, *p* < .001, *β* = 0.830). Including a stimulation variable in the model did not yield a significant improvement (*F*_(1,13778)_ = 0.745, *p* = 0.388), suggesting that the strong negative correlation between theta power and negative rule prediction errors held under both anodal and sham conditions.

**Figure 6.**
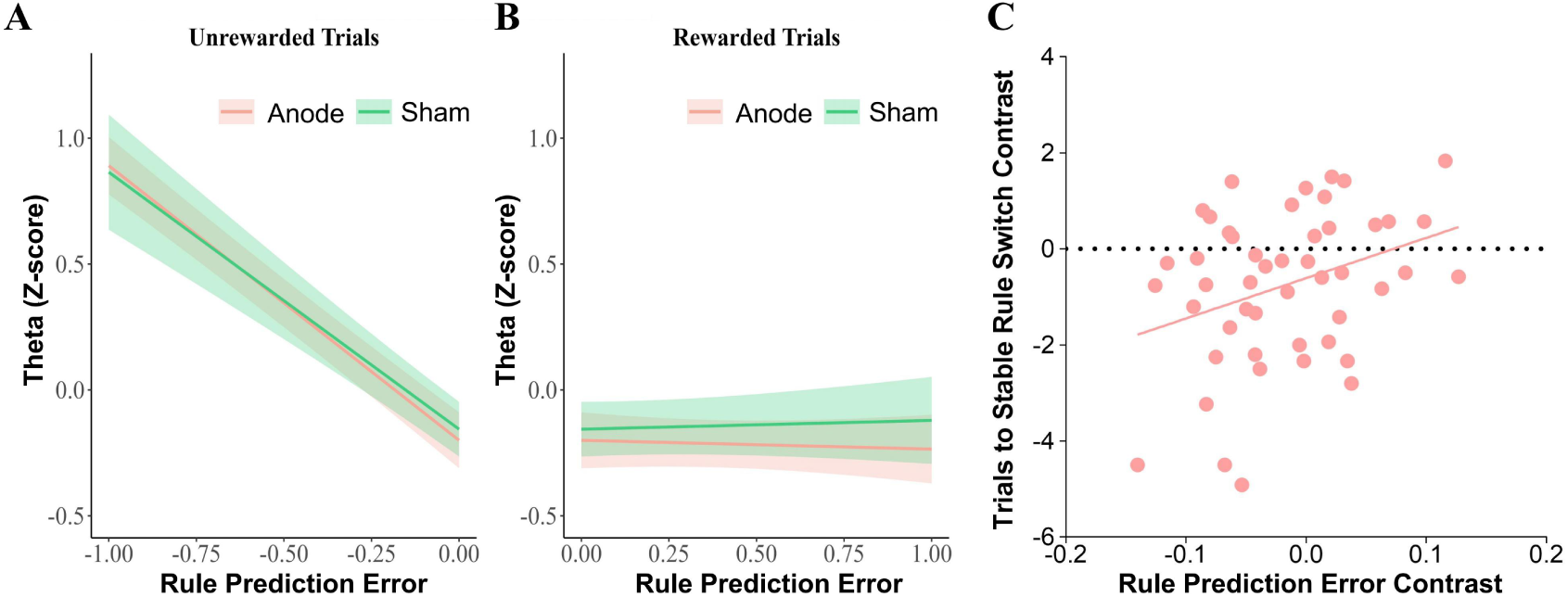
Results of the regression model between theta power and rule prediction errors across stimulation conditions. (A) Unrewarded trials: theta power increases as rule prediction errors decrease in both anodal and sham conditions. (B) Rewarded trials: theta power remains relatively stable across rule prediction errors values in both conditions. Shaded areas indicate 95% confidence intervals (CIs). Lines represent the trial-by-trial relationship between estimated rule prediction errors and the theta power extracted from the clusters shown in Figure 5. (C) The figure shows the relationship between the rule prediction error contrast (sham - anode) under negative feedback and the contrast in trials to achieve a stable rule switch. A positive correlation was observed (r = 0.330, p = 0.022), indicating that a greater reduction in rule prediction error following anodal stimulation was associated with a larger increase in the number of trials required to reach stable switching behavior.

### Anodal tDCS Reduced Sensitivity to Negative Feedback Predicts Delayed Rule Switching

To examine whether the reduced sensitivity to negative feedback under anodal stimulation influenced behavioral adaptation, we conducted an across-subject correlation analysis between the rule prediction error contrast (sham - anode) under negative feedback and the number of trials to stable rule switch contrast (sham - anode). A significant positive correlation was found (*r* = 0.330, *p* = 0.022, Figure 6C), suggesting that participants with a greater reduction in the absolute value of negative rule prediction error following anodal stimulation (i.e. less sensitive to negative feedback) also required more trials to adjust to the new rule. This finding supports the idea that attenuated neural sensitivity to negative feedback may impair timely behavioral adjustment.

### Anodal tDCS Increases Frontal Theta Power During Baseline Period

Previous research has demonstrated that activity in the MFC at 500 ms before stimulus onset is also critical for rule-switching tasks (Cole et al., 2024). We averaged theta power within the 500 ms interval during the baseline period before stimulus onset separately for the anodal and sham conditions (Figure 7A and Figure 7B) and conducted a topographical comparison by subtracting anode from sham. We found that theta power in the 500 ms preceding stimulus onset was significantly higher under anodal stimulation compared to sham. The difference was localized to frontal electrodes, with significantly elevated theta power observed at FCz, FC1, FC2, Fz, F1, F2, AFz, AF3, and AF4 (Figure 7C). Individual-level analysis within the predefined frontal ROI (theta band, -500 to 0 ms) revealed a significant difference, as shown in Figure 7D. (*t*_(47)_ = -2.53, *p* = 0.0148, Cohen’d = -0.365). This pattern suggests that anodal stimulation modulates theta activity differently across task phases—enhancing it during the baseline period and reducing it following feedback. These phase-specific effects are expected, as participants are presented with outcome-related feedback during the feedback period, and the blunted theta activity after anodal stimulation is closely linked to the processing of feedback events.

**Figure 7.**
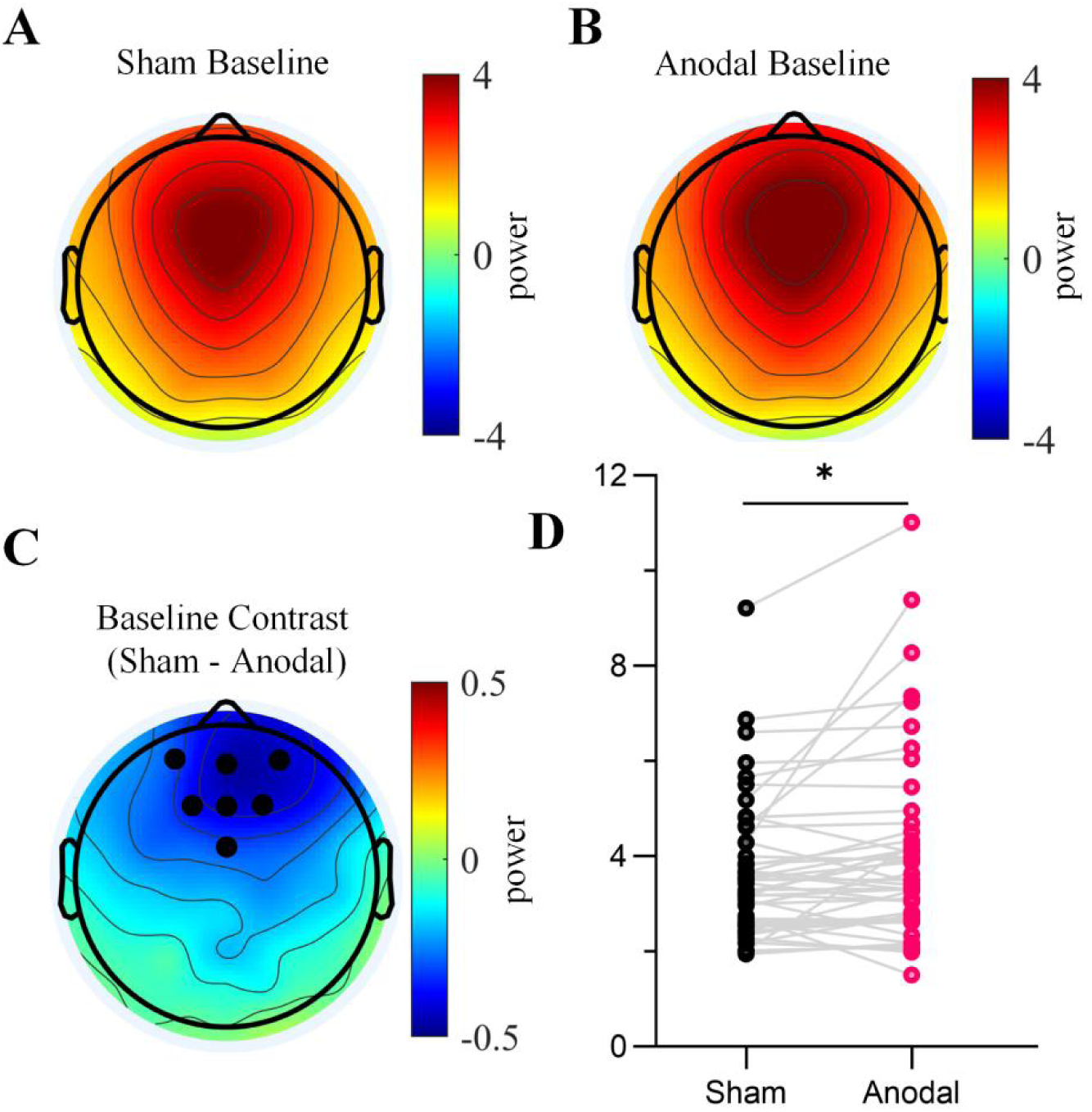
Topographical distribution of theta power during the baseline period. (A) Topography of theta power under sham stimulation. (B) Topography of theta power under anodal stimulation. (C) Difference map showing the contrast between anodal and sham conditions. (D) Results of a paired-sample t-test comparing average theta power across ROI channels between the two stimulation conditions.

### Theta Reduction Tied to Rule Switching Deficits

To examine how pre-stimulus baseline activity and attenuated post-feedback theta power collectively impair cognitive flexibility, we extracted the contrast between theta power from feedback phase and from the -500 to- 0 ms interval from pre-stimulus baseline period. We conducted a general linear model (GLM) at the across-subject level using changes in theta power from feedback phase and baseline period between sham and anodal conditions and related these neural measures to the behavioral measure of trials to stable rule switch contrast (Table 1) Across subjects, the predictor *Theta Neg-Pos contrast* from feedback was negatively associated with trials to stable rule switch (*β* = −2.136, *SE* = 0.926, *t* = −2.31, *p* = 0.026), whereas *Baseline contrast* did not reach statistical significance (*β* = −0.345, *SE* = 0.209, *t* = −1.65, *p* = 0.105). We found that changes in theta power after feedback under anodal stimulation, relative to sham, significantly predicted changes in trials to stable rule switch. Greater reductions in theta power under anodal stimulation, compared to sham, predicted diminished stable rule switching performance, as reflected by larger absolute contrast values. These findings indicate that a more blunted theta response in response to errors correlated with poorer performance in trials to stable rule switch.

**Table 1.** GLM: Stable Rule Switch contrast ∼ Theta Neg-Pos contrast + Baseline contrast.

| Predictor | Estimate | Std. Error | t-value | p-value |
| --- | --- | --- | --- | --- |
| Intercept | -0.754 | 0.219 | -3.44 | 0.001 |
| Theta Neg-Pos<br>contrast | -2.136 | 0.926 | -2.31 | 0.026 * |
| Baseline contrast | -0.345 | 0.209 | -1.65 | 0.105 |

### Causal Pathway from Frontal Theta to Adaptive Rule Switching Mediated by Prediction Error under Anodal Stimulation

Anodal stimulation was found to influence theta power, rule prediction error, and trials to stable rule switch. We next attempted to clarify the relation between theta power, rule prediction error, and trials to stable rule switch by fitting mediation models under anodal and sham stimulation using trial-level data. For the sham condition (Figure 8A), the path from (feedback-locked) theta power to rule prediction error (*a* path) was significant (*β* =–0.024, *SE* = 0.006, *z* =–3.715, *p* < 0.001), indicating that higher theta power was associated with lower rule prediction error. Specifically, increased theta was associated with lower signed (negative) rule prediction error, entailing an association between increased theta and increased absolute rule prediction error. However, the path from rule prediction error to trials to stable rule switch (*b* path) was non-significant (*β* = 0.015, *SE* = 0.328, *z* = 0.045, *p* = 0.964), as was the direct path from theta power to trials to stable rule switch (c′ path; *β* =–0.035, *SE* = 0.040, *z* =–0.881, *p* = 0.378). Accordingly, the indirect effect (*a* × *b*) was not significant (*β* =–0.000, *SE* = 0.008, *z* =–0.043, *p* = 0.966, Cohen’s *d* = -0.0062), nor was the total effect (*β* =–0.036, *SE* = 0.039, *z* =–0.906, *p* = 0.365). These results suggest that rule prediction error does not mediate the relationship between theta power and trials to stable rule switch under sham stimulation.

**Figure 8.**
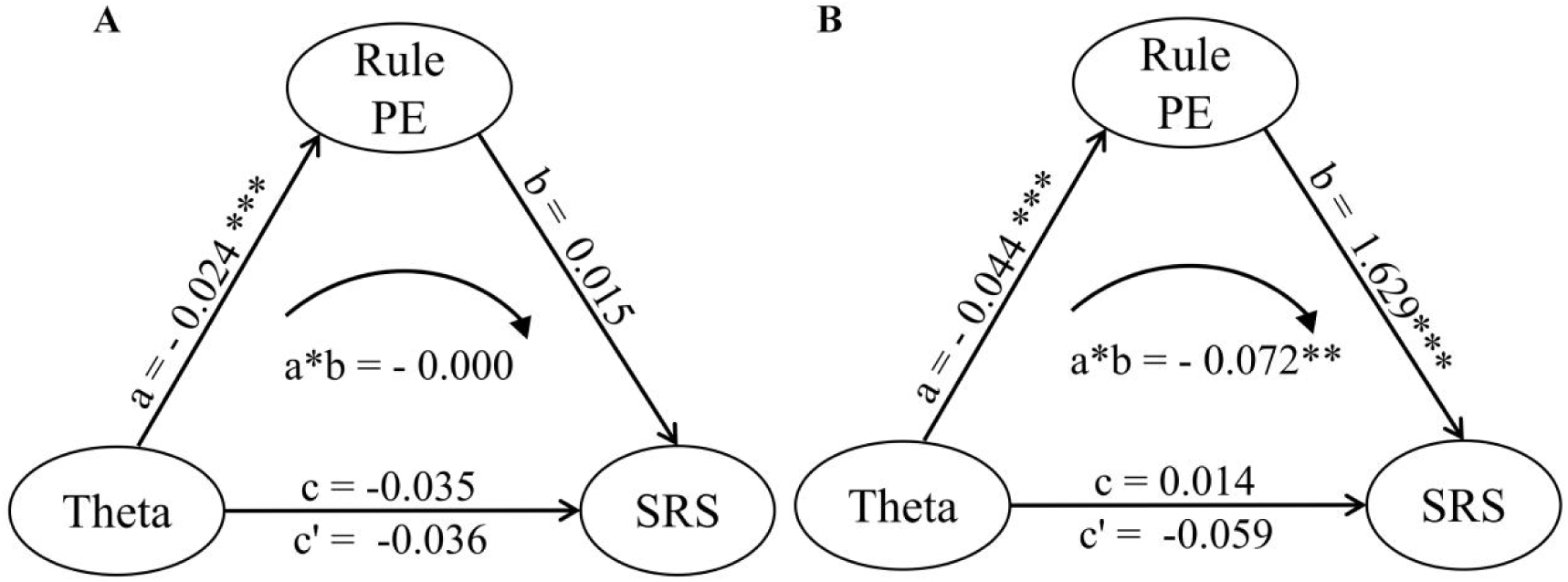
Mediation effect of theta power on trials to stable rule switch under sham (A) and anodal (B) stimulation. (A) Under the sham condition, a significant negative association was observed between theta power and rule prediction error. The indirect effect (a × b) was not significant, nor was the total effect. (B) The mediation analysis revealed a significant indirect pathway by which theta power influenced behavioral adaptation (Stable Rule Switch) via its effect on rule prediction error. The indirect effect was significant while the direct effect of theta on Stable Rule Switch was non-significant, suggesting full mediation. *p < 0.05; **p < 0.01; ***p < 0.001.

We next conducted the same mediation analysis under anodal stimulation (Figure 8B). Theta power significantly predicted rule prediction error (*a* path; *β* =–0.044, *SE* = 0.006, *z* =–6.941, *p* < 0.001), indicating that reduced theta power was associated with increased rule prediction error: Again, increased theta was associated with lower signed (negative) rule prediction error, so increased theta was associated with increased absolute rule prediction error. Rule prediction error, in turn, significantly predicted trials to stable rule switch (*b* path; *β* = 1.629, *SE* = 0.500, *z* = 3.258, *p* = 0.001) (similarly, blunted rule predicted error increased trials to stable switch). The direct effect of theta power on trials to stable rule switch (*c*′ path) was not significant (*β* = 0.014, *SE* = 0.039, *z* = 0.355, *p* = 0.723). Importantly, the indirect effect (*a* × *b*) was significant (*β* = –0.072, *SE* = 0.025, *z* =–2.909, *p* = 0.004, Cohen’s *d =* -0.420), indicating a reliable mediation of theta power on behavior via its influence on rule prediction error. The total effect (*c* path) of theta on trials stable rule switch did not reach significance (*β* =–0.059, *SE* = 0.040, *z* =–1.482, *p* = 0.138), suggesting that the behavioral impact of theta is primarily indirect. These findings support the hypothesis that theta oscillations contribute to adaptive rule updating by modulating the processing of rule prediction errors.

These results suggest that theta oscillations alone are not sufficient to drive behavioral adaptation unless they modulate the computation of rule prediction errors. Compared to the sham condition, in which no significant mediation effect was observed, the anodal condition exhibited theta suppression that led to reduced rule prediction error and altered behavioral adaptation. These results reinforce the mediating role of prediction error in linking frontal theta dynamics to adaptive rule switching.

## Discussion

In this study, we employed high-definition transcranial direct current stimulation as a tool to explore causal relationships between neural oscillatory activity and cognitive flexibility, specifically focusing on theta oscillation during a probabilistic reversal learning task. Our results show that after anodal tDCS, theta power at negative feedback decreased compared to the sham stimulation. This change in brain activity appears to be causally linked to a behavioral shift, as participants under anodal stimulation showed blunted rule (but not stimulus-action) prediction errors and required more trials to stabilize on the new rule following a reversal.

Our hypotheses were guided by the Sync model of cognitive control (Verbeke and Verguts, 2019), which holds that strategy switches (e.g., in which task set is currently active) are signaled via rule prediction errors and subsequent theta power increase. Behaviorally, this model fit the best, and the anodal stimulation jointly influenced theta power, rule prediction error, and trials to rule switch, as this model predicts. Anodal stimulation may interfere with the functional role of theta oscillations in transmitting prediction error signals critical for updating latent rule representations (Senoussi et al., 2022).

We did not observe significant differences in overall accuracy or response time between stimulation conditions, while stimulation robustly modulated more rule-switching-sensitive measures such as trials to stable rule switch. This dissociation suggests that conventional behavioral metrics may not be sufficiently sensitive to detect changes in internal cognitive dynamics, especially when task performance remains at ceiling levels. In contrast, trials to stable rule switch captures trial-by-trial adaptation patterns that reflect how participants internalize and respond to rule uncertainty, making it a more sensitive index of neuromodulatory effects on cognitive flexibility. Similar dissociations have been reported in prior tDCS and neuropharmacological studies (Leite et al., 2013; Albein-Urios et al., 2019; Panitz et al., 2022), where neural or computational markers showed significant modulation in the absence of overt changes at accuracy.

The reduction in rule prediction error sensitivity and the increased trial number to reach stability after rule reversal may indicate that anodal tDCS does not uniformly facilitate cognitive flexibility. Studies have shown that the effects of anodal tDCS on executive function tasks typically depend on task complexity and cognitive load (Fertonani and Miniussi, 2017). In low cognitive load tasks, anodal tDCS may enhance cortical activity and improve performance. However, in high cognitive load tasks, its effects may be limited or even counterproductive, potentially leading to task interference due to excessive excitation (Iuculano and Cohen Kadosh, 2013; Hill et al., 2016). This may explain why anodal tDCS did not enhance cognitive flexibility in our study. While anodal tDCS is generally considered to enhance cortical excitability, its effects are subject to multiple interacting factors.

Theta-band activity in the MFC, particularly in response to negative outcomes, has been consistently implicated in the signaling of cognitive conflict and adaptive control (Cavanagh and Frank, 2014; Liu et al., 2022). The decreased theta power in the negative-positive feedback period after anodal stimulation may reflect an underlying neural mechanism related to diminished feedback processing and learning(Cavanagh and Frank, 2014; Senoussi et al., 2022). The observed decrease in theta power could represent a reduction in the engagement of MFC involved in evaluating and updating rule contingency (Cavanagh et al., 2012; Cohen and Donner, 2013). Interestingly, this feedback-phase effect contrasts with the increased theta activity observed during the pre-stimulus period (baseline period) under anodal stimulation. Such opposite-phase effects are reminiscent of dopaminergic modulation, where tonic increases in dopamine are thought to compress the dynamic range of phasic responses, effectively blunting prediction error signaling (Frank, 2005). Analogously, anodal stimulation may elevate baseline theta activity, which in turn reduces the brain’s capacity to generate strong phasic responses to salient events such as feedback. This mechanism could underlie the reduced sensitivity to feedback and impaired adaptation observed under anodal stimulation.

Although both anodal and cathodal tDCS over the MFC reduced frontal theta power, only anodal stimulation led to a diminished absolute rule prediction error during negative feedback. Anodal stimulation may disrupt the precise temporal coordination of theta bursts necessary for the rule prediction error computation, thereby weakening the fidelity of prediction error encoding and reducing the subject’s responsiveness to environmental changes. In contrast, cathodal stimulation, while also reducing theta power, did not alter prediction error estimates. This indicates that the suppression of oscillatory power per se is not sufficient to disrupt higher-order learning processes. Instead, cathodal tDCS may exert a more generalized inhibitory effect on cortical excitability without targeting the specific mechanisms involved in feedback-driven updating. This polarity-specific effect underscores the importance of considering not only the magnitude but also the functional significance of stimulation-induced neural changes. The selective effect of anodal stimultion aligns with prior findings showing that anodal tDCS often produces more robust or selective effects on behavior and neural dynamics compared to cathodal stimulation (Jacobson et al., 2012; Dedoncker et al., 2016; Reinhart et al., 2017). The absence of cathodal effects here is therefore not surprising, but rather consistent with this broader literature. Taken together, these results highlight a dissociation between neural signatures (theta power) and their computational consequences (prediction error updating) and point to anodal tDCS as an effective means of perturbing the MFC circuitry involved in adaptive learning.

In conclusion, by combining these neural and behavioral observations within a cognitive model-based framework, we establish a causal link between theta power modulation and altered cognitive flexibility, with rule prediction error serving as a key computational mechanism that mediates this relationship. Our study highlights the potential of tDCS in combination with computational modeling as a tool for investigating causal relationships in cognitive neuroscience, particularly regarding how brain oscillations influence prediction error signaling and, in turn, rule adaptation. Future research could explore whether these effects are modulated by individual differences in baseline cognitive flexibility or how long-term exposure to tDCS may influence these learning dynamics.

## Supplementary Information

## Supplementary Note 1

Goodness-of-fit indices (Mean LL, AIC, and wAIC) for the three computational models (RW, ALR, and Sync) across different stimulation conditions (sham, anode, cathode) (Table 1). Higher log-likelihood (LL) and wAIC values, and lower AIC values, reflect better model fit. Results suggested that across all stimulation conditions, the Sync model consistently outperformed the RW and ALR models, demonstrating the best overall fit to the behavioral data.

**Table 1.** Goodness-of-fit measures.

| Model | Conditions | Mean | SD | Mean | SD | Mean | SD |
| --- | --- | --- | --- | --- | --- | --- | --- |
|  |  | LL | LL | AIC | AIC | wAIC | wAIC |
| RW | Sham | -113.34 | 18.55 | 230.67 | 37.10 | 0.01 | 0.04 |
|  | Anode | -110.35 | 17.94 | 224.79 | 35.87 | 0.08 | 0.25 |
|  | Cathode | -113.99 | 18.91 | 231.99 | 37.81 | 0.01 | 0.05 |
| ALR | Sham | -111.84 | 19.06 | 229.67 | 38.12 | 0.03 | 0.10 |
|  | Anode | -112.97 | 18.46 | 231.94 | 36.92 | 0.09 | 0.28 |
|  | Cathode | -112.51 | 19.00 | 231.03 | 38.00 | 0.04 | 0.18 |
| Sync | Sham | -95.53 | 20.96 | 199.06 | 41.93 | 0.97 | 0.13 |
|  | Anode | -94.16 | 19.58 | 196.31 | 39.16 | 0.84 | 0.36 |
|  | Cathode | -96.79 | 19.93 | 201.59 | 39.85 | 0.95 | 0.19 |

<sup>a</sup> Results of LL, AIC, and wAIC computations over subjects are shown for each of three models. For LL and wAIC, high values indicate a better fit, whereas for AIC a low value indicates a good fit.

## Supplementary Note 2

Perseverative errors were defined as the number of errors from each rule reversal until the participant’s first successful switch to the correct rule. To assess whether stimulation conditions influenced perseverative errors, we conducted a repeated measures ANOVA. The results showed no significant differences in perseverative errors across the three stimulation conditions (Figure 2A) (*F*_(2, 93.98)_ = 0.51, *p* = 0.60; sham: *M* = 4.13, *SD* = 1.07; anode: *M* = 4.34, *SD* = 1.06; cathode: *M* = 4.23, *SD* = 1.19). This indicates that perseverative errors were comparable across conditions, suggesting that the stimulation did not significantly alter participants’ initial tendency to persist with the previous rule before successfully switching.

We assessed regression errors. Participants in the anode stimulation group showed significantly more regression errors compared to the sham group, indicating a stronger tendency to persist with the previous rule despite changes in task contingencies (Figure 2B) (*F*_(1.88, 88.45)_ = 7.23, *p* = 0.0082; sham: *M* = 0.57, *SD* = 0.78; anode: *M* = 1.17, *SD* = 1.21; cathode: *M* = 0.94, *SD* = 1.02). This suggests greater difficulty in maintaining the correct rule following reversals under anode stimulation.

**Supplementary Figure 2:**
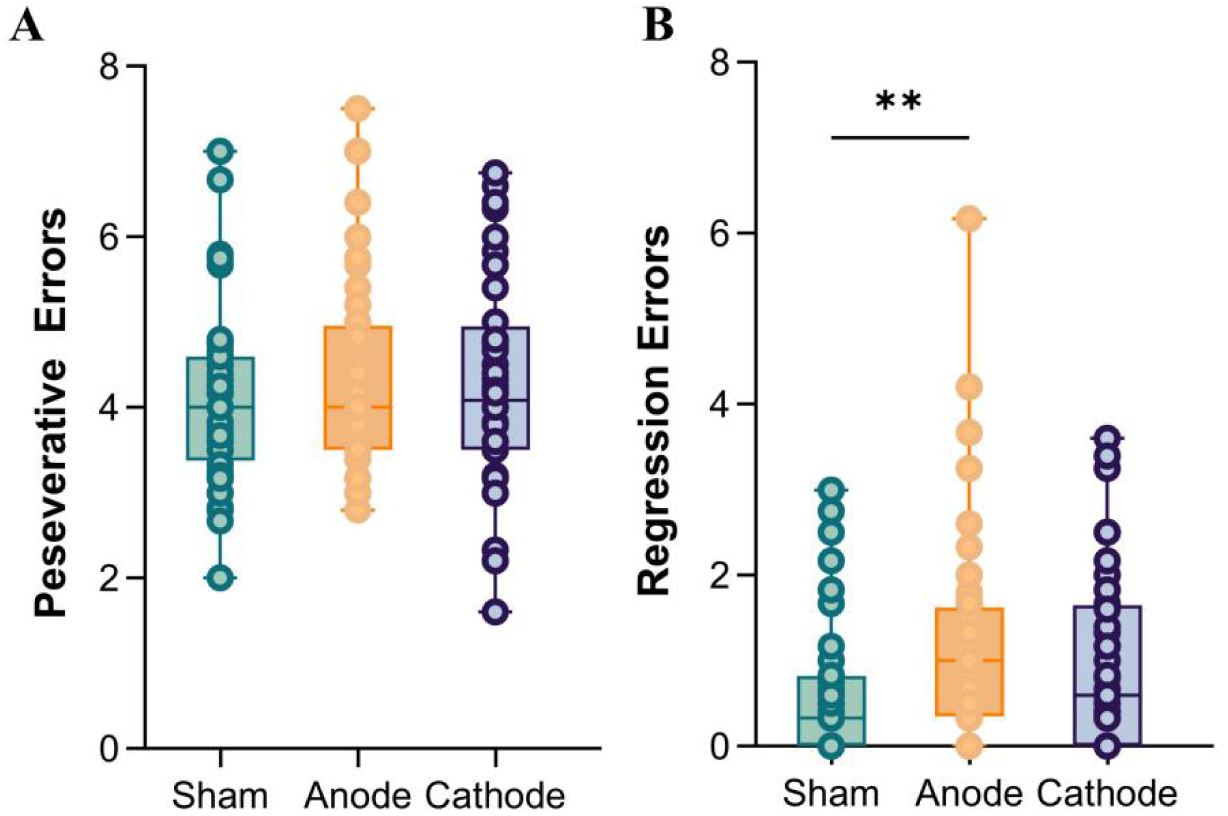
Behavioral results of perseverative errors and regression errors across stimulation conditions. Figure 2. (A) Perseverative errors: No significant differences were observed in perseverative errors across the sham, anodal, and cathodal conditions. (B) Regression errors: participants in the anodal condition exhibited significantly more regression errors compared to the sham condition (*p* < 0.01). No significant differences were found between the cathodal and sham conditions. Error bars represent the standard error of the mean (SEM).

## Supplementary Note 3

In the main text, we reported that only the rule prediction error associated with rule switching showed significant differences across the three stimulation conditions. Additionally, we compared other model parameters, including mapping learning rate, switch learning rate, cumulation, and temperature, but found no significant differences among the three stimulation conditions.

**Supplementary Figure 3:**
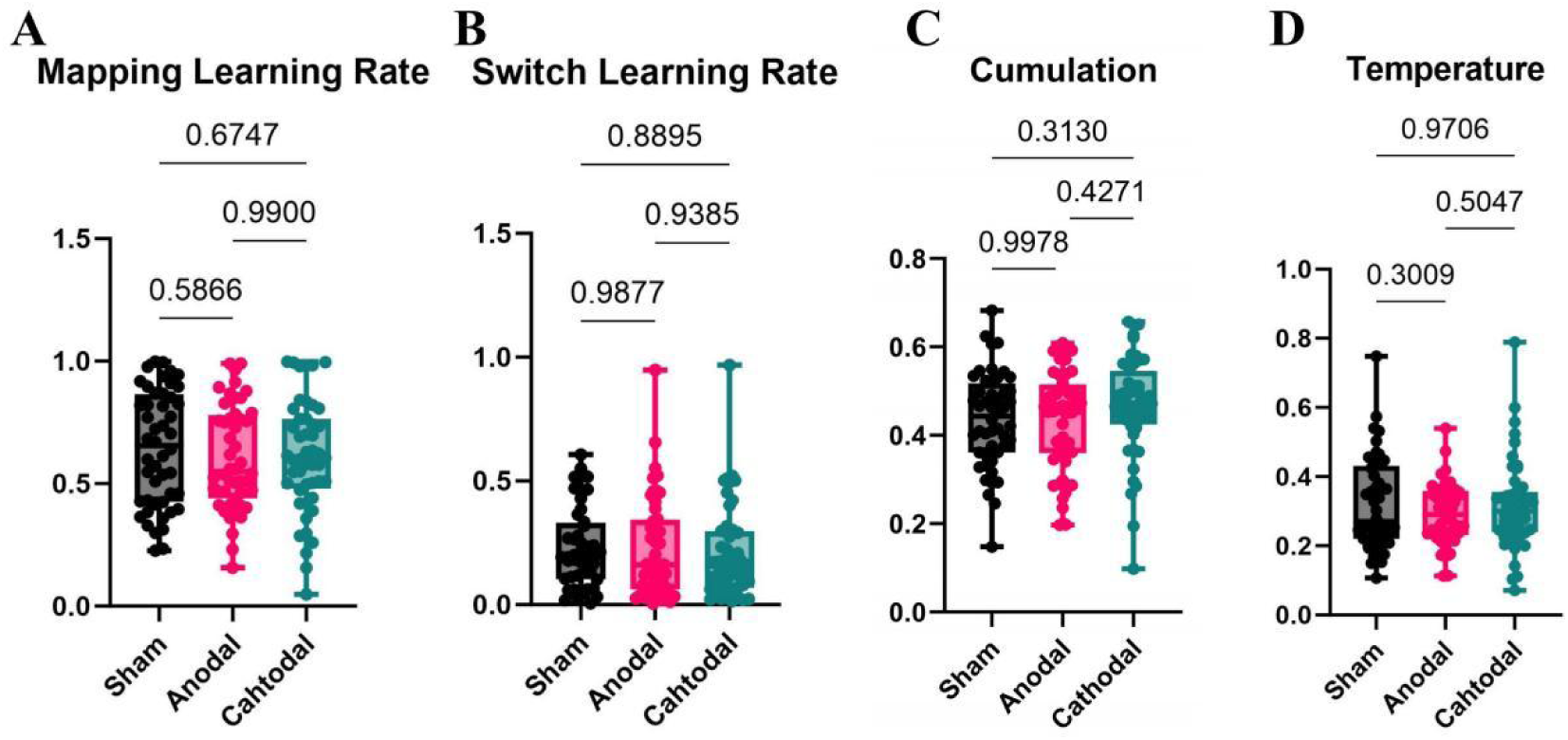
Model Parameters comparison Figure 3. Comparison of model parameters across the three stimulation conditions (anode, sham, and cathode) using repeated measures ANOVA. (A) Mapping Learning Rate: Represents the parameter governing how the model updates rule mapping. (B) Switch Learning Rate: Indicates how quickly the model adapts to rule changes. (C) Cumulation: Reflects the extent to which rule prediction errors accumulate over time. (D) Temperature: Represents the degree of randomness in decision-making. For all parameters (A-D), there were no significant differences across the three stimulation conditions (anode, sham, and cathode).

## Supplementary Note 4

We conducted an additional control analysis focusing specifically on negative low-level prediction errors, defined as stimulus-action prediction errors. A linear mixed-effects model was applied, with stimulation condition as a fixed effect and subject as a random intercept. This analysis revealed no significant effect of stimulation on negative stimulus–action prediction errors (*F*_(1, 6834)_ = 0.476, *p* = 0.621). These results indicate that anodal stimulation did not significantly influence low-level prediction error processing, suggesting that its effect was selective to rule-level prediction errors.

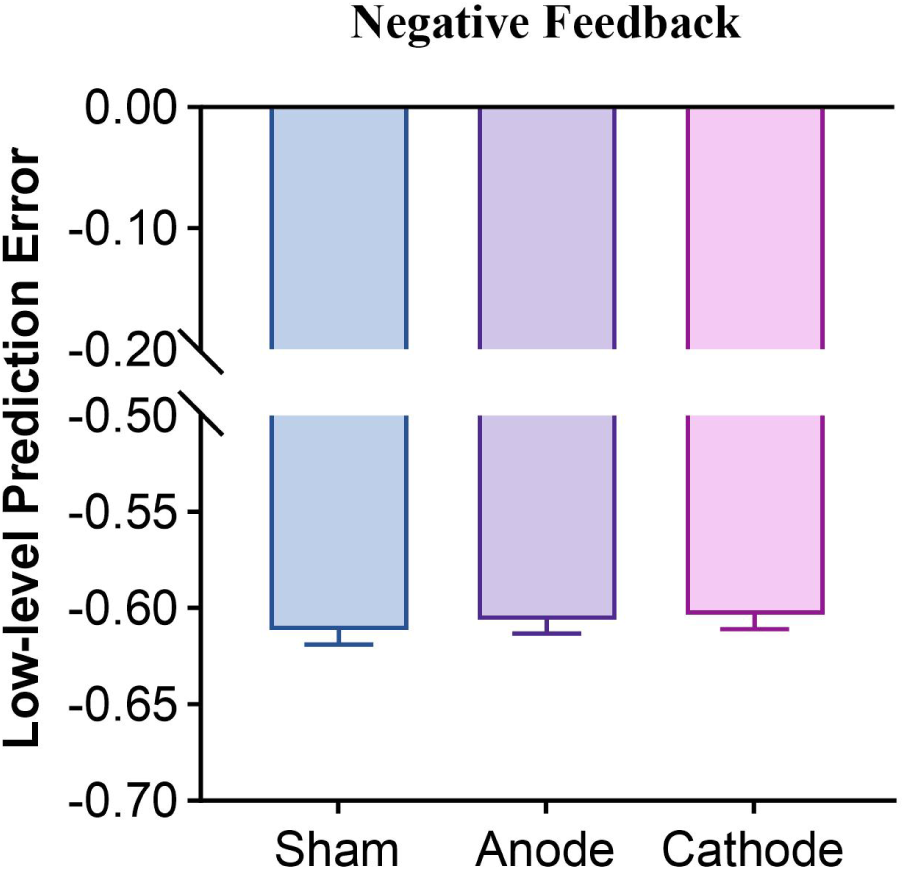
Figure 4. Low-level prediction error under negative feedback across stimulation conditions. Bars represent the mean signed prediction error values for sham, anodal, and cathodal stimulation. No statistically significant differences emerged between conditions. Error bars indicate the standard error of the mean (SEM). A broken y-axis is used to improve visualization.

## Supplementary Note 5

To examine whether cathodal stimulation also modulates theta activity in the probabilistic reversal learning task, we conducted the same analyses as for the sham and anodal conditions. Specifically, we performed: (1) time-frequency analysis to identify significant theta clusters in the cathodal condition for the Neg-Pos feedback contrast, (2) a comparison of theta clusters between sham and cathodal conditions, and (3) an analysis of theta power differences between sham and cathodal conditions during the baseline period (500 ms prior to stimulus onset).

As shown in Supplementary Figure 5A, a significant theta cluster was identified in the cathodal condition, occurring approximately between 0 and 600 ms. Figure 5B presents the topography of this theta cluster, with significant electrode sites marked in black. Figure 5C illustrates the significant clusters in the contrast between sham (Neg-Pos) and cathodal (Neg-Pos) conditions, highlighting a decrease in theta activity following cathodal stimulation.

Additionally, Figure 5D examines theta power during the 500 ms baseline period preceding stimulus onset. Importantly, Supplementary Figure 5D illustrates that no significant difference was observed between the sham and cathodal conditions during the same pre-stimulus interval (*t*_(47)_ = -1.64, *p* = 0.108). The top-left panel of Figure 5D shows the sham condition, while the bottom-left panel presents the cathodal condition, with the difference topography displayed on the right. No significant electrode sites were identified, indicating that theta power did not differ significantly between sham and cathodal conditions during the baseline period.

**Supplementary Figure 5:**
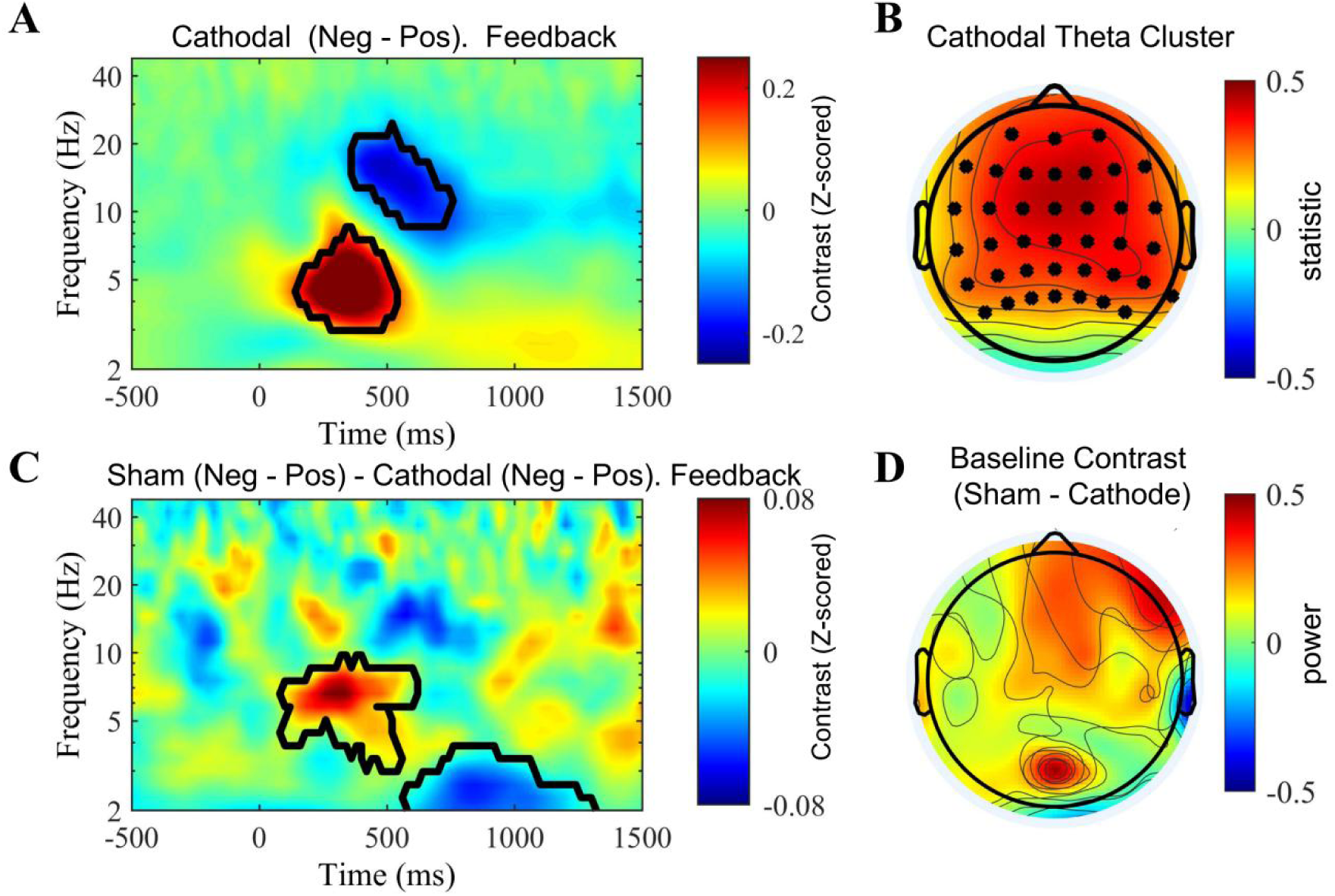
Time-frequency analysis of cathodal condition Figure 5: (A). Time-frequency analysis of Neg-Pos feedback contrast under cathodal stimulation. B. Topographic map of the theta cluster during the cathodal condition. (C) Significant clusters in the contrast between sham (Neg-Pos) and cathodal (Neg-Pos) conditions, showing differences between sham and cathodal stimulation. (D) Topographic maps for sham and cathodal conditions 500 ms before stimulus onset. The rightmost map shows the baseline contrast (500 ms before stimulus onset) between cathodal and sham conditions.

## Supplementary Note 6

We also provide an analysis of the relationship between rule prediction error and theta power under the cathodal stimulation condition. This analysis follows the same approach as the one used for the sham and anodal conditions in the main text.

We observed that, similar to the other two stimulation conditions, theta power and rule prediction error exhibited a negative correlation under cathodal stimulation in the negative feedback condition. However, no such relationship was found in the positive feedback condition.

**Supplementary Figure 6:**
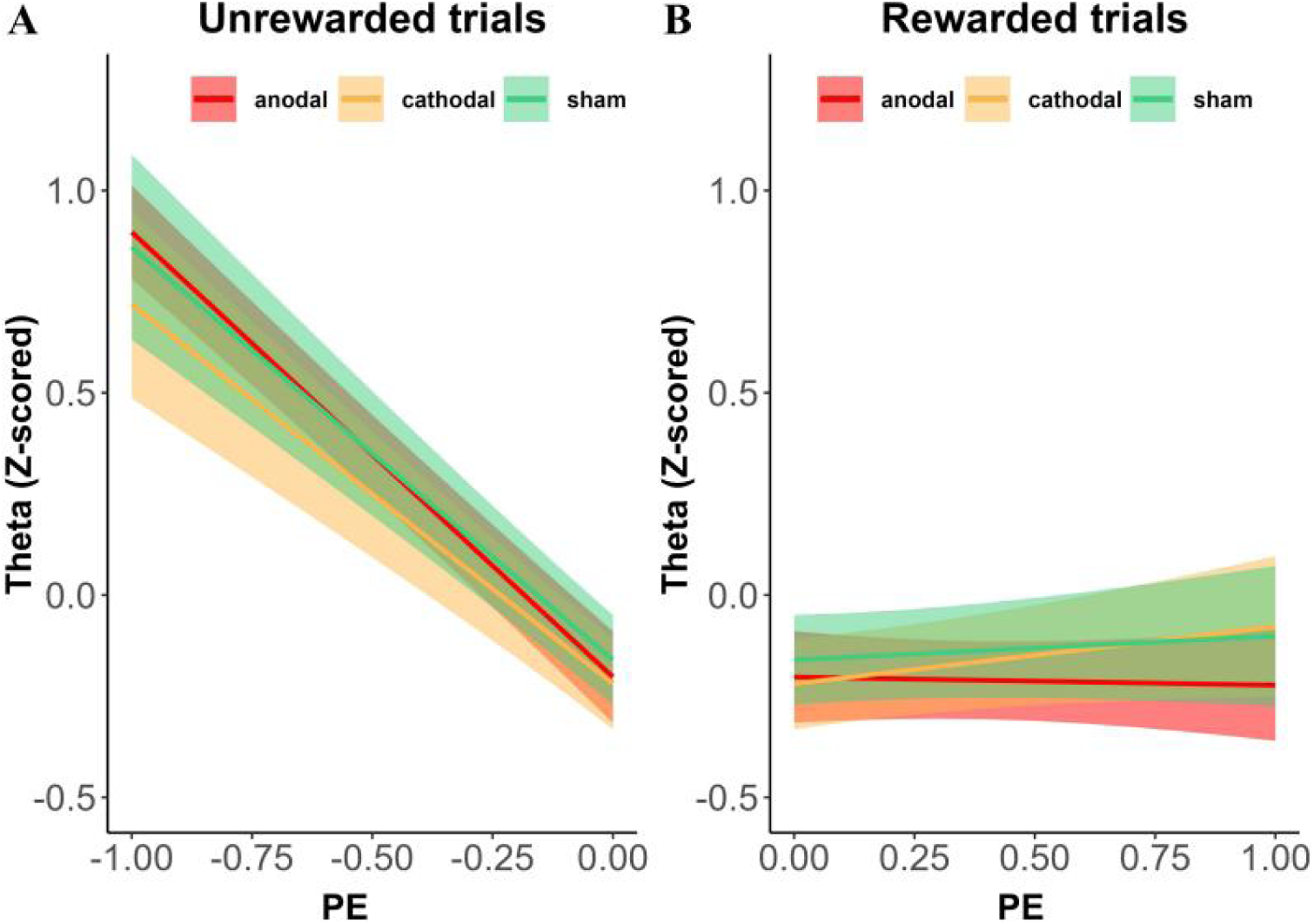
Relationship between Rule Prediction Error and Theta Figure 6. Results of linear regression between theta power and rule prediction errors across stimulation conditions (sham, anode and cathode). The plot illustrates a negative linear relationship between theta power and negative rule prediction error under anode and sham stimulation. Lines represent the trial-by-trial relationship between estimated rule prediction error and the mean theta power extracted from the clusters shown in Figure 5. Shaded areas indicate 95% confidence intervals (CIs).

## Supplementary Note 7

This figure illustrates the distribution of the Neg - Pos theta contrast (sham minus anode) across participants, extracted from the significant group-level cluster in the Neg–Pos feedback comparison. Each dot represents one participant’s z-scored theta difference, with the boxplot indicating the mean and standard error. A one-sample t-test revealed that the effect was significant (t(47) = *t*_(47)_ = 6.03, *p* < .001, *M* = 0.209, *SD* = 0.241), confirming that, on average, theta power was reduced following anodal stimulation relative to sham. Notably, 40 out of 48 participants (approximately 83.3%) consistent with this overall pattern, suggesting that the suppressive effect of anodal stimulation on feedback-related theta activity was highly consistent across individuals.

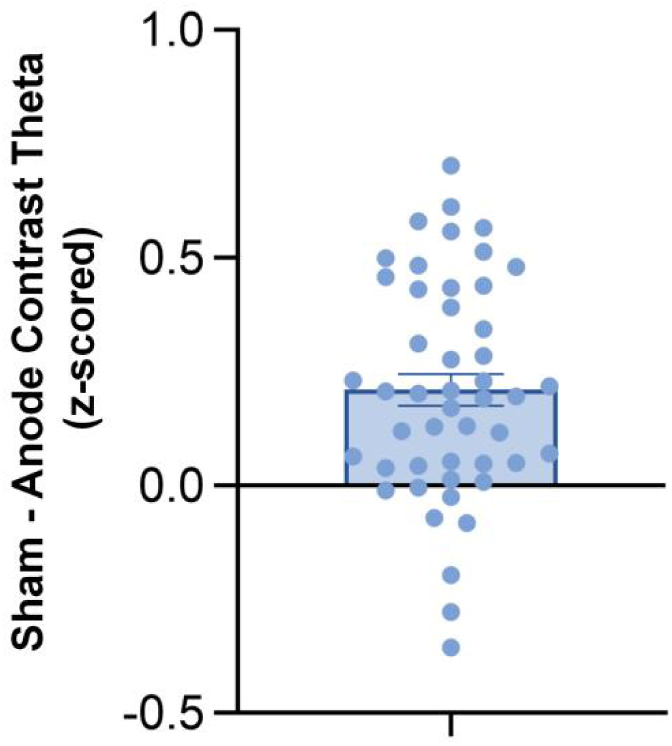
Figure 7. Individual-level distribution of feedback-related theta modulation (sham-anode contrast)

## Supplementary Note 8

To explore whether theta oscillations mediate the influence of rule prediction error on behavioral adaptation under cathodal stimulation (Supplementary Figure 8), we conducted a mediation analysis using single-trial data from the rule reversal event. Theta power significantly predicted prediction error (a path: *β* = 0.030, *SE* = 0.007, *z* = 4.128, *p* < 0.001), suggesting that higher theta power was associated with larger prediction errors. However, the path from prediction error to Stable Rule Switch (b path: *β* = 0.166, *SE* = 0.477, *z* = 0.349, *p* = 0.727) and the direct path from theta to Stable Rule Switch (c′ path: *β* =–0.028, *SE* = 0.050, *z* =–0.562, *p* = 0.574) were both non-significant. Correspondingly, the indirect effect did not reach significance (*β* = 0.005, *SE* = 0.015, *z* = 0.340, *p* = 0.734), and the total effect of theta on Stable Rule Switch was also non-significant (*β* =–0.033, *SE* = 0.046, *z* =–0.716, *p* = 0.474). These results suggest no evidence of a mediating role of theta power under cathodal stimulation. Interestingly, although theta power significantly decreased under cathodal stimulation, this neural change did not translate into altered behavioral performance. The mediation analysis revealed that rule prediction error remained unchanged, and neither the indirect nor direct effects of theta on Stable Rule Switch reached significance.

**Supplementary Figure 8.**
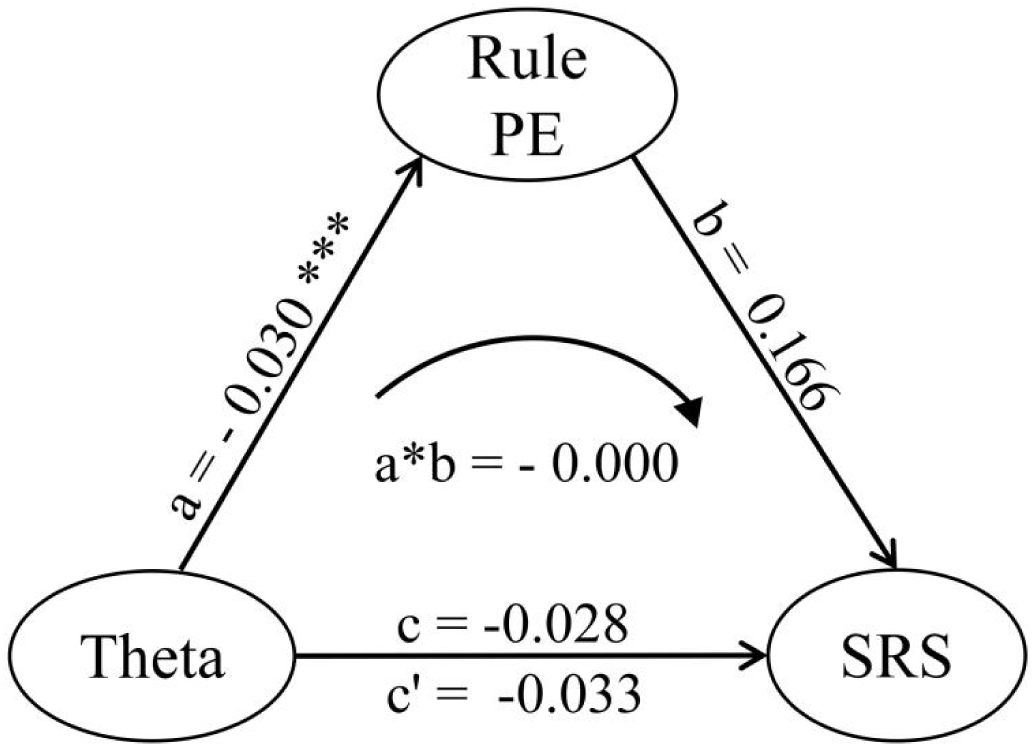
Cathodal Mediation Analysis. Figure 8. Result of mediation analysis under cathodal condition. Both the indirect and total effects are not significant.

